# Microvascular structure variability explains variance in fMRI functional connectivity

**DOI:** 10.1101/2024.03.22.585979

**Authors:** François Gaudreault, Michèle Desjardins

## Abstract

The influence of regional brain vasculature on resting-state fMRI BOLD signals is well documented. However, the role of brain vasculature is often overlooked in functional connectivity research. In the present report, utilizing publicly available whole-brain vasculature data in the mouse, we investigate the relationship between functional connectivity and brain vasculature. This is done by assessing interregional variations in vasculature through a novel metric termed vascular similarity. First, we identify features to describe the regional vasculature. Then, we employ multiple linear regression models to predict functional connectivity, incorporating vascular similarity alongside metrics from structural connectivity and spatial topology. Our findings reveal a significant correlation between functional connectivity strength and regional vasculature similarity, especially in anesthetized mice. We also show that multiple linear regression models of functional connectivity using standard predictors are improved by including vascular similarity. We perform this analysis at the cerebrum and whole-brain levels using data from both male and female mice. Our findings regarding the relation between functional connectivity and the underlying vascular anatomy may enhance our understanding of functional connectivity based on fMRI and provide insights into its disruption in neurological disorders.

## 1 Introduction

Understanding how behavior emerges from the underlying anatomy is one of the main goals of neuroscience. A starting point in this quest is to explore how brain function is related to its underlying anatomy. In recent years, this has been done by extensively studying the relationship between functional connectivity (FC) and structural connectivity (SC). FC is the statistical relationship between specific neurophysiological signals in time across different regions of the brain and SC represents the anatomical connections between those regions.

In humans, studying this relationship at the whole-brain scale has been facilitated by advancements in magnetic resonance imaging (MRI) technology. The assessment of FC at this scale was enabled through the development of functional MRI (fMRI) (Belliveau et al. 1991; Ogawa et al. 1992; Bandettini et al. 1992; Kwong et al. 1992) while diffusion-weighted MRI allowed the visualization of nerve tracts (Howe et al. 1992; Pierpaoli et al. 1996) for SC. The combination of these imaging techniques permitted the comparison and prediction of resting-state FC with tract-tracing data (Koch et al. 2002; Greicius et al. 2009; Honey et al. 2009; Damoiseaux and Greicius 2009). In fixed mouse brains, SC can be studied through injections of viral vectors to trace individual axonal projections between regions. This method led to the mesoscale structural connectome in the mouse (Oh et al. 2014), which has become a standard for comparing SC and FC (Grandjean et al. 2017; Coletta et al. 2020; Mahani et al. 2023).

Nevertheless, in mammals, the most common technique for measuring large-scale FC is still fMRI which is not a direct measurement of neuronal activity but rather a vascular correlate. Indeed, neuronal activity leads to an increase in oxygen consumption and cerebral blood flow which results in a net positive fMRI signal: the blood-oxygen-level-dependent (BOLD) signal (Ogawa et al. 1990; Attwell and Iadecola 2002; Buxton 2013). However, it is well known that this signal depends upon the underlying brain vasculature (Chen et al. 2010; Vigneau-Roy et al. 2014; Drew 2019; Tsvetanov et al. 2021). Considering this, should models using SC alone be sufficient to describe fMRI-based FC?

To address this question, we leveraged whole-brain vasculature datasets recently made available (Quintana et al. 2019; Todorov et al. 2020; Kirst et al. 2020; Ji et al. 2021; Wu et al. 2022). Specifically, the data from Ji et al. (2021) is utilized to quantify the relationship between regional vascular variability and FC. We accomplished this by using a new metric called vascular similarity (VS) in multivariable linear regression models, which have been used extensively in the study of FC and SC (Goñi et al. 2014; Vázquez-Rodríguez et al. 2019; Zamani Esfahlani et al. 2022). This kind of model allows us to quantify the impact of regional vascular variability in a functional connectivity model using other common predictors derived from spatial topology and SC.

## 2 Materials and methods

### 2.1 Functional connectivity data

#### 2.1.1 fMRI data

In this article, three openly available resting-state fMRI datasets are used : a dataset of N = 10 anesthetized female C57BL/6J mice^1^ We obtained the raw data which we processed and used for computing functional connectivity. Both datasets in anesthetized mice were acquired using a combination of medetomidine and isoflurane anesthesia (0.05 mg/kg bolus and 0.1 mg/kg/h IV infusion, plus 0.5% isoflurane) and artificial ventilation at a rate of 90 BPM. The mice of the awake male dataset were headfixed, an habituation protocol was followed 10-15 days after the headpost surgery (Gutierrez-Barragan et al. 2022). In both male datasets, the resting-state fMRI scans were acquired using a single-shot echo planar imaging (EPI) sequence on a 7.0 Tesla animal scanner with the following parameters: TR/TE=1000/15 ms, flip angle=60^°^, matrix=100×100, 18 coronal slices (voxel size 0.23 × 0.23 × 0.60 mm^3^) and 1920 time points. In the female dataset, the scans were acquired on an 11.7 Tesla Bruker animal scanner using gradient echo EPI with the following parameters: TR/TE=1200/15 ms, flip angle=60^°^, matrix=90×60, 28 coronal slices (voxel size 0.18 × 0.15 × 0.40 mm^3^) and 180 time points. Female mice underwent three scans per session across two sessions, resulting in a total of six scans per mouse and an aggregate of 60 scans. Also, this dataset contained T2-weighted structural images of each mouse at both imaging sessions which were used as anatomical templates.

#### 2.1.2 Preprocessing and confound correction

We preprocessed all datasets using the open-source RABIES software^2^ (Desrosiers-Grégoire et al. 2024). Briefly, for each subject, a mean volumetric EPI image was derived using a trimmed mean across all frames after an initial motion realignment. This mean EPI image allows the estimation of head motion by registering each EPI frame to it using rigid transformations. For the female dataset, the structural T2-weighted images were initially corrected for inhomogeneities (Wang et al. 2017) and then used to generate an unbiased structural template using a re-implementation of the ANTs template construction pipeline^3^ (Avants et al. 2011). This newly generated unbiased template was then itself registered, using a nonlinear registration, to the DSURQE ex-vivo T2 MRI template (Dorr et al. 2008; Richards et al. 2011; Steadman et al. 2014; Ullmann et al. 2013) which had been registered to the Allen Institute for Brain Science (AIBS) Common Coordinate Framework space version 3^4^. After this, each EPI image was also corrected for inhomogeneities and slice timing (slice timing correction was made using AFNI’s 3dTshift) (Cox 1996) and registered to the same session anatomical scan. Then all the transformations required for these corrections were concatenated to allow the resampling of each EPI frame to generate EPI time series in the reference atlas space. For the male dataset, the same steps were followed with the exception that the EPI scans were used to generate the unbiased template which was registered to the DSURQE atlas instead of (unavailable) structural scans.

For consistency, the confound correction applied to the EPI time series was the same for the male and female anesthetized datasets. First, frames with prominent corruption were censored as in Power et al. (2012). Framewise displacement was measured across time and each frame surpassing 0.05 mm of motion, together with 1 backward and 2 forward frames, were removed. Voxelwise detrending was applied to remove first-order drifts. Using ordinary least squares, the 6 rigid motion parameters and the mean signal from the white matter and cerebrospinal fluid masks were modeled at each voxel and regressed from the data. To normalize variance, each image was separately scaled according to its total variance. Finally, a spatial Gaussian smoothing filter was applied at 0.3 mm full-width at half maximum (FWHM).

Additionally, to reduce the motion artifacts in the awake dataset, the motion sources were automatically removed using a modified version of the ICA-AROMA classifier (Pruim et al. 2015), where classifier parameters and anatomical masks are instead adapted for rodent images.

#### 2.1.3 FC computation

Following the confound correction for each subject, the mean fMRI timecourse of each region was extracted. The correlation between the timecourses of each pair of regions was calculated, yielding a functional connectivity matrix for each mouse. The subject-specific matrices were then Fisher z-transformed before being averaged over subjects to obtain a group FC matrix. The final matrix results from averaging the FC matrices of both hemispheres. The brains were parcelled according to a mid-ontology level of the AIBS Common Coordinate Framework version 3 (CCFv3). The anatomical regions that were considered are the 213 anatomical regions of Oh et al. (2014). The two missing regions are due to the removal of the subdivision of the subiculum into its dorsal and ventral part from the Common Coordinate Framework version 2 (CCFv2) to the CCFv3 as well as the distinction between the subiculum and the prosubiculum (Wang et al. 2020). FC matrices of two different subsets of the 211 regions of interest (ROIs) were analyzed. The first subset was an FC matrix of size 78 × 78 composed of the same subset of regions as in Mills et al. (2018) minus the subdivision of the subiculum: the 78 regions in this subset are all part of the cerebrum. The other subset was composed of the 200 ROIs that were present in all subjects after registration in the two anesthetized datasets. This led to an FC matrix of size 200 × 200 (to obtain the full list of 200 regions see supplementary material). We will differentiate the use of the FC matrix of 78 and 200 regions by referring to them as the cerebrum and whole-brain FC respectively. In the male awake dataset the subparaventricular zone was excluded in the whole-brain FC because it was not present in all the subjects. This leads to 199 ROIs instead of 200, but to avoid confusion when referring to the whole-brain regions set we will not make the distinction.

### 2.2 Structural connectivity data

For structural connectivity data, the matrix from Knox et al. (2018) was used. It was generated using a mathematical model at the level of 100 *µ*m voxels using 428 of the 491 anterograde tracing experiments in wild-type C57BL/6J mice, mapping fluorescently labeled neuronal projections brain-wide from the AIBS connectivity atlas^5^ (Oh et al. 2014). This model of SC was chosen because it offers improved predictions at the regional level over the model from Oh et al. (2014). The weights of the SC matrix are the normalized connection strengths (Knox et al. 2018). This matrix is directed as opposed to the FC, hence to compare the two the SC matrix must be undirected. This was accomplished by averaging the directed SC matrix with its transpose. Moreover, the structural connectivity matrix wasn’t directly used in the linear model: instead, the communicability of the undirected SC was used. This was done to better explain the observed patterns of FC, as it is known that low SC strength doesn’t imply low FC strength (Damoiseaux and Greicius 2009). Communicability is a metric that describes the efficiency with which information can be transmitted across a network using direct and indirect pathways. The definition of communicability used is given by Crofts and Higham (2009). We began by normalizing each connection weight of the network. We divided them by the product 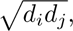 where *d_i_* is the generalized degree of the node *i* to obtain a new matrix *A^′^*. The communicability is then given by the matrix exponential

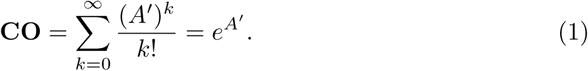

The matrix exponential computes the weighted sum of walks of all lengths between pairs of nodes while penalizing longer walks by the 1*/k*! coefficients. Ultimately, pairs of nodes with stronger and more direct connections should have higher communicability weight. The steps to obtain the structural connectivity matrices are illustrated in figure 2B.

### 2.3 Brain vasculature data

To compare functional connectivity with regional vascular anatomy, vascular graphs of three adult male mice from Ji et al. (2021) were used. In their paper, whole-brain microvasculature was imaged in fixed brains to the level of capillaries with serial fluorescence two-photon microscopy and modeled into a vascular graph. In this representation, edges represent vessels and nodes represent branching points. The available data was the position of centerline voxels composing edges, the voxels composing nodes, and the list of nodes connecting individual edges. Each brain was also registered to the Allen Brain Atlas template, making it possible to label each voxel to a brain region. Using the nodes composing a region of interest, we built a regional graph of the vasculature from which the regional features were extracted. This process was repeated for all ROIs in each brain. The procedure to retrieve each regional graph’s features is illustrated in figure 1.

**Fig. 1.**
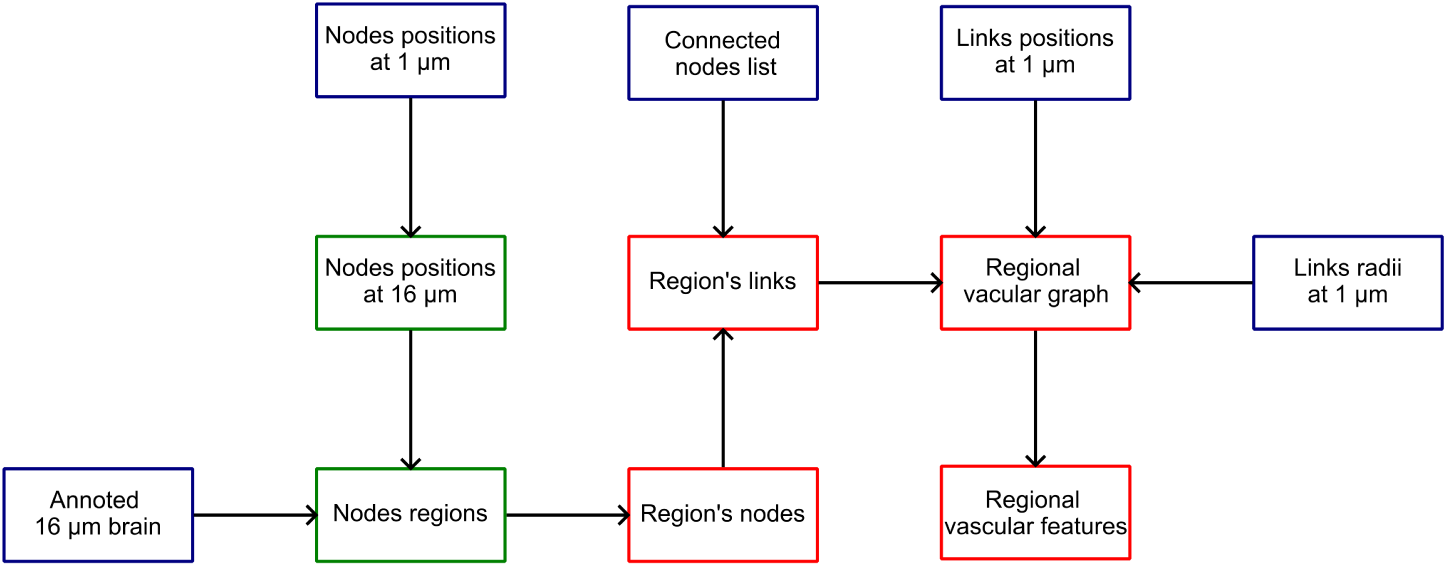
Numerical processing of the whole-brain vasculature graphs from Ji et al. (2021). Each box represents an object and the arrows signify a function that transforms the source object into the destination object. The blue boxes denote the objects constituting the comprehensive brain graph, the green boxes correspond to transformed objects at the whole-brain level, and the red boxes represent objects specific to distinct brain regions.

The regional features were then averaged over the two hemispheres and the three subjects. Eight vascular features were tested in the anesthetized datasets: average vessel tortuosity, average capillary length and radius, proportion of length covered by non-capillary vessels, proportion of volume filled by vessels, and length, branching points and vessel density. Capillaries were defined as vessels with a median radius of their skeleton voxel radii lower than 3.5 *µ*m as in Ji et al. (2021). Using the regional features a VS matrix is built using equation 2

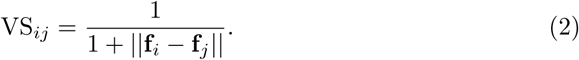

where **f***_i_* is the feature vector of the region *i* (as illustrated in figure 2A) and ||**f***_i_* − **f***_j_*|| is the Euclidean distance between the feature vectors. To ensure equivalent scaling, each feature was z-scored across all regions before computing the values for VS.

**Fig. 2.**
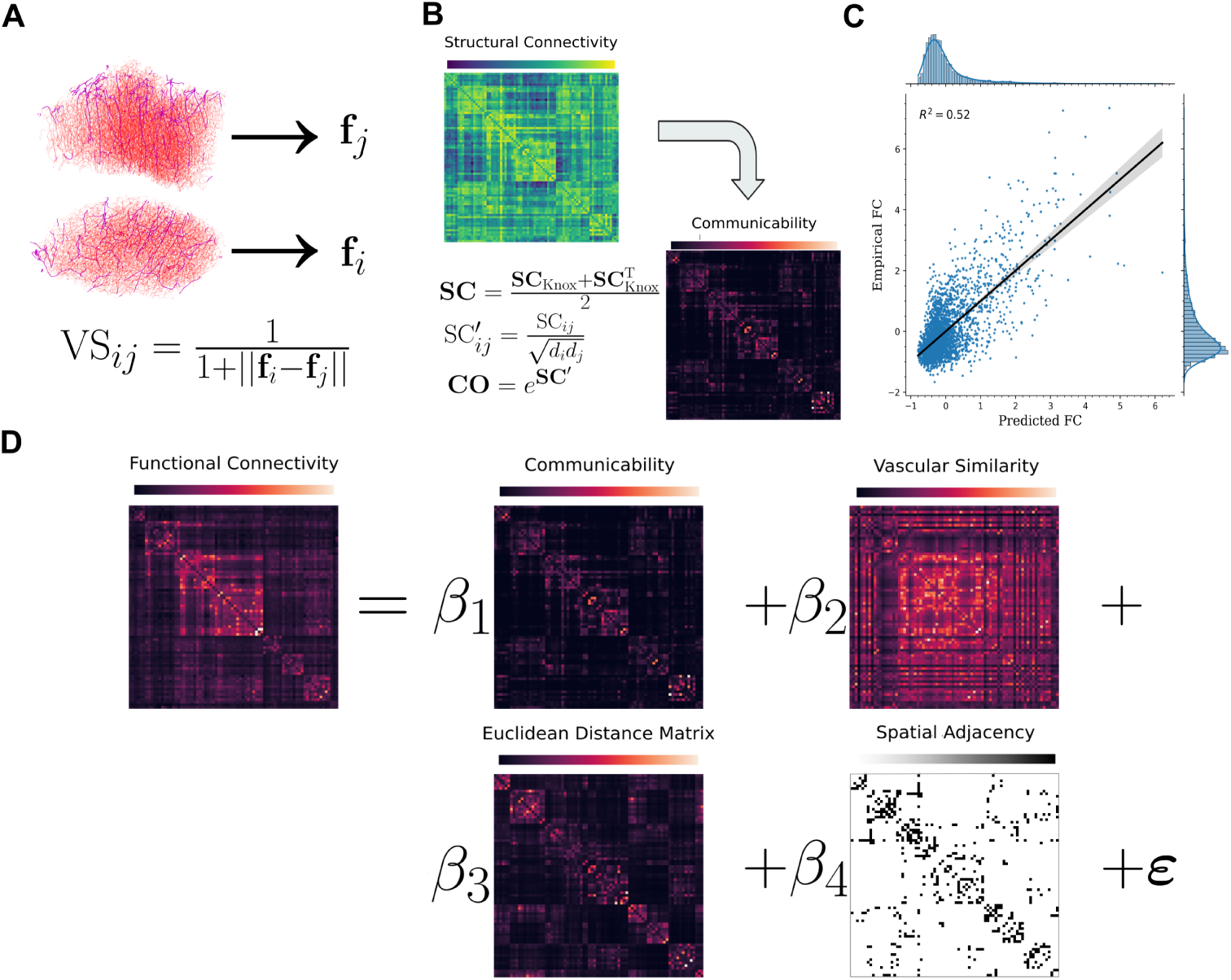
The full functional connectivity model. A) Representation of the vascular similarity strength between two vascular regional graphs *i* and *j*. B) Procedure for deriving the communicability predictor from the SC matrix. C) The full FC model of the female dataset using the subset of 78 ROIs. Empirical and predicted FC represent z-scored functional connectivity strength between regions. D) Illustration of the model, *β* parameters are estimated using ordinary least squares. For visualization purposes, the regions are ordered by FC communities computed using the Louvain method for community detection.

Several of these characteristics may not significantly contribute to the explained variance of functional connectivity (as expressed by the coefficient of determination). To explore which vasculature features are the most relevant to describe FC, VS matrices were made using only one feature. Then, simple linear models of FC were built using only the single-feature VS as an independent variable. The features that were ultimately used for building the VS of the full multiple linear regression model were the four features that had on average the highest coefficient of determination (*R*^2^) in the single-feature simple linear regression FC models. We chose to pick four features to mitigate the effect of collinearity between them. In practice, when more of the eight original features are used, their addition has been found to reduce correlation values.

As a means to determine the statistical significance of the correlation between functional connectivity and vascular similarity, two null models were used. The first one is called the random model and consists of making a new VS matrix by randomly shuffling the vector of features of each region before calculating VS. The second is labeled the spatial autocorrelation preserving model. As the name implies, this null model keeps the original spatial autocorrelation of each feature before calculating the VS. This process was accomplished through the application of BrainSMASH (Burt et al. 2020), a method that resamples the values of each vascular feature while maintaining the original spatial autocorrelation by leveraging the regional distance matrix. For both of these models, 1000 different VS matrices were computed. Subsequently, Pearson correlation coefficients between the FC matrices and the null VS matrices values for each region pair were computed.

### 2.4 Multiple linear regression models

To better understand the global association between functional and vascular similarity, several multiple linear models were built using combinations of the following independent variables: vascular similarity, communicability, inverse of the Euclidean distance between the geometric center of the regions, and spatial adjacency. The latter is a binary metric with a value of 1 if two regions are adjacent in space and 0 if they aren’t. When these four predictors were used in a model we call it the full functional connectivity model. These predictors are illustrated in figure 2D. FC matrices and independent variables were z-scored before calculating the model parameters and only values below the diagonal of the matrices of the matrices were used for the linear models. The coefficients of the linear models were found using ordinary least squares. In the linear models employed in this study, intercept terms were computed. However, for the purposes of this analysis, these values were excluded from detailed discussion, as they were found to be negligible.

We compared the results from our anesthetized FC models using the 78 × 78 matrices to those built in Mills et al. (2018) which we will call the Mills models. In the Mills models, the 80 × 80 FC matrix was constructed using resting-state fMRI data from male subjects under isoflurane anesthesia, and was parcelled in accordance with CCFv2. The SC matrix is the inter-region connectivity model from Oh et al. (2014). The predictors derived from the structural connectivity were the communicability and the matching index. The model also makes use of gene expression measured with in situ hybridization from AIBS as described in Lein et al. (2007). A subset of regional gene expression energies was then z-scored across all regions and Pearson’s correlations were computed between region pairs and Fisher z-transformed, which is called the correlated gene expression (CGE) in the Mills model. To facilitate comparative analysis, we also computed CGE as described in Mills et al. (2018) and included it in our model.

Finally, the multivariable linear regression models using the 200 × 200 anesthetized FC matrices were calculated to examine the relationship between VS and FC at the whole-brain scale. The awake dataset was also used to assess the impact of anesthesia on the model’s results. Subsequently, to investigate the overlap between the vascular similarity and functional connectivity, binary FC and VS matrices were made by thresholding and binarizing the group female FC matrix and the VS matrix from 30% to 10% of connection density with 1% interval as in Mills et al. (2018). Then the intersection between the binary VS and FC matrices of the same density was calculated. This resulted in 21 overlap matrices; the edges that were present in more than half of these matrices were kept to produce an average binary overlap matrix.

### 2.5 Alignment of vascular features with cortical hierarchical organization

Finally, we investigated the relationship between vascular features and the hierarchy of functional organization of the mouse isocortex. It has been shown that in the human brain, many structural and functional properties align with a hierarchy of increasing functional integration (Mesulam 1998; Huntenburg et al. 2018). The same conclusion was also drawn for the mouse brain in Fulcher et al. (2019). In the latter, the ratio of T1-weighted to T2-weighted (T1w:T2w) images is used as a marker of hierarchical specialization. To quantify the alignment of the vascular features with the functional hierarchy, we began by performing principal component analysis (PCA) on the four vascular features used to construct the vascular similarity matrices. Next, we computed the Spearman correlation coefficient across regions between the first principal component and the T1:T2 ratio reported in Fulcher et al. (2019). Out of the 40 isocortex regions studied in their article, the vascular features were not available in two of them, specifically the posterior auditory area and the primary somatosensory area, unassigned. The correlations values were computed with the 38 remaining regions. Furthermore, we also computed the Spearman correlation coefficient of the first principal component with the consensus gradient of Fulcher et al. (2019). This gradient orders areas along a putative functional hierarchy, from primary somatosensory to integrative prefrontal areas.

All analyses were conducted using Python. Notably, the Allen SDK library was used to access the AIBS datasets and the Graph-tool library was utilized during vascular graph analysis.

## 3 Results

### 3.1 Relationship between functional connectivity and single-feature vascular similarity

The values of the coefficient of determination for simple linear regression models between FC and single-feature VS are presented in table 1. The explained variance by single-feature VS is higher for all features with the 78 regions models compared to the 200 regions ones. The explained variance difference between males and females was less than 1% for most features except for the average capillary length. The correlation between the anesthetized male and female datasets group functional connectivity was computed, revealing significant correlations of (r=0.816) for cerebrum FC and (r=0.668) for whole-brain FC. For both sexes and region sets, the worst coefficients of determination are the proportion of the vessel length covered by non-capillary vessels and the proportion of the regional volume filled by vessels. The four features that explain the most variance on average are the branching point density (BPD), the average capillary length, tortuosity, and radius. They also correspond to the combination of features that explain the most variance in all the simple linear regression FC-VS models. These features are mostly driven by the capillary network of the region.

**Table 1.**
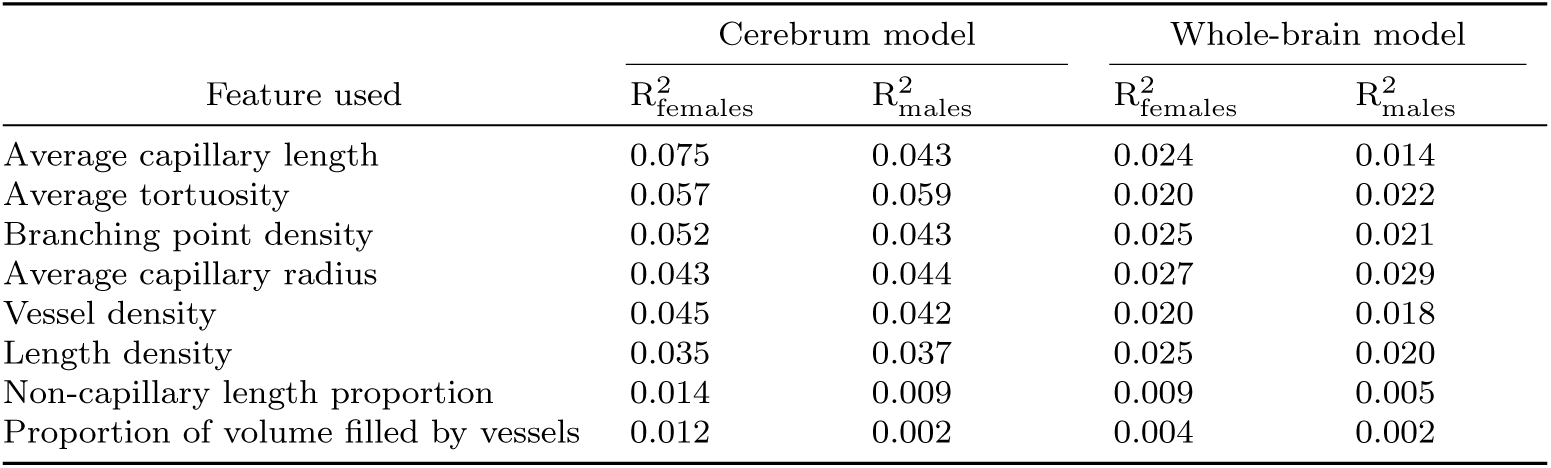
Explained variance of functional connectivity by the different single-feature vascular similarity as the sole predictor in the linear model across both female and male datasets using the sets of regions of the cerebrum and the whole brain.

We computed the correlation between each pair of vascular features. Among the above four, the highest pairwise correlation values for the cerebrum are the BPD and the average capillary length (r=0.593). For the whole brain, it’s the BPD and tortuosity pair (r=0.515). Across all features, the highest correlation values are between the BPD and the vessel density for cerebrum and whole-brain models (r=0.988 in females, r=0.959 in males). This is expected because most vessels split or recombine into two. High correlation values were also found for BPD and the length density (r=0.927 in females, r=0.830 in males).

### 3.2 Comparison with the linear model of Mills and alignment of vascular features with the functional hierarchy

Functional connectivity models of the cerebrum were built using the four-feature vascular similarity and compared to the values of Mills FC models. These results are presented in table 2. At this scale, vascular similarity alone explains about 15.9% and 15.1% of the variance in female and male FC. This is significant because these values are much higher than the average determination coefficient obtained using the spatial autocorrelation preserving null models ⟨R^2^⟩=0.03 (SD=0.02) and ⟨R^2^⟩=0.02 (SD=0.02) for female and male datasets (see supp. figure 2 for the correlation values distribution of the null models for the cerebrum). When VS is made using all eight features the explained variance diminishes to 12.1% and 11.0% for female and male FC. It can also be observed that VS increases the explained variance of FC when added to the communicability, Euclidean distance and spatial adjacency model. Furthermore, in the full model, the VS has the second highest standardized *β* coefficient after communicability despite explaining less variance than spatial adjacency and Euclidean distance using simple linear regression. In the cerebrum, VS is also less correlated to the inverse of the Euclidean distance than to all the other predictors as can be seen in supp. figure 1. For the full list of standardized parameters of the cerebrum model of this study see supp. table 1.

**Table 2.** Comparison between the anesthetized FC models of Mills and the cerebrum FC models of this study. The R^2^ values for some of the Mills models included are calculated using the matching index as an additional predictor. These models are denoted by the addition of an extra term (**MI**). The *β_i_* are the standardized coefficients of each parameter. **F**^^^**C** is the predicted FC connection weights. **CO** is the communicability connection weights. **CGE** is the correlation of selected genes expressions between each pair of ROIs. **VS** is the Euclidean distance similarity measure between the vascular feature vectors of each pair of ROIs. In our models, **ED** represents the inverse of one plus the Euclidean distance between the ROIs centroids. In the Mills models, these Euclidean distances are log-transformed instead. **SA** is the spatial adjacency between each pair of ROIs, if two regions are adjacent the value of the pair is 1, 0 otherwise.

| Linear model | $R^2_{\text{females}}$ | $R^2_{\text{males}}$ | $R^2_{\text{Mills}}$ |
| --- | --- | --- | --- |
| $\hat{\mathbf{F}}\mathbf{C} = \beta_1 \mathbf{CO}$ | 0.416 | 0.411 | 0.227 |
| $\hat{\mathbf{F}}\mathbf{C} = \beta_1 \mathbf{VS}$ | 0.159 | 0.151 | - |
| $\hat{\mathbf{F}}\mathbf{C} = \beta_1 \mathbf{CGE}$ | 0.310 | 0.308 | 0.412 |
| $\hat{\mathbf{F}}\mathbf{C} = \beta_1 \mathbf{ED}$ | 0.265 | 0.290 | - |
| $\hat{\mathbf{F}}\mathbf{C} = \beta_1 \mathbf{SA}$ | 0.302 | 0.330 | - |
| $\hat{\mathbf{F}}\mathbf{C} = \beta_1 \mathbf{ED} + \beta_2 \mathbf{SA}$ | 0.355 | 0.388 | 0.549 |
| $\hat{\mathbf{F}}\mathbf{C} = \beta_1 \mathbf{CO} + \beta_2 \mathbf{VS}$ | 0.479 | 0.470 | - |
| $\hat{\mathbf{F}}\mathbf{C} = \beta_1 \mathbf{CO} + \beta_2 \mathbf{ED} + \beta_3 \mathbf{SA} (+\beta_4 \mathbf{MI})$ | 0.464 | 0.478 | 0.584 |
| $\hat{\mathbf{F}}\mathbf{C} = \beta_1 \mathbf{CO} + \beta_2 \mathbf{VS} + \beta_3 \mathbf{ED} + \beta_4 \mathbf{SA}$ | 0.520 | 0.529 | - |
| $\hat{\mathbf{F}}\mathbf{C} = \beta_1 \mathbf{CO} + \beta_2 \mathbf{CGE} + \beta_3 \mathbf{ED} + \beta_4 \mathbf{SA} (+\beta_5 \mathbf{MI})$ | 0.493 | 0.502 | 0.615 |

As for the comparison between the models in this study and the Mills models, the communicability alone explained significantly more variance in our models than in the Mills models. This can be attributed to the use of the Knox model of SC. In fact, when using the SC matrix from Oh et al. (2014), as in Mills’s article, to derive the communicability matrix, an important decrease in the explained variance is observed in both females and males (17.5% and 15.9%). Besides the communicability model of FC, the R^2^ values of Mills were higher in all models. Moreover, spatial models of FC based on the Euclidean distance and spatial adjacency explain significantly less variance than the Mills model. One other considerable difference between the models of this study and the one from Mills (Mills et al. 2018) is how the Euclidean distance is used. In the model of Mills, the Euclidean distance is log-transformed before being used. In our cerebrum spatial models, if the log-transformed Euclidean distance is utilized, a decrease in explained variance is observed in the female and male (R^2^=0.347 and R^2^=0.369).

When considering the four predictors models, the model of female and male FC with the highest explained variance is the one that includes the VS rather than the CGE. These models still explain lower variance than the Mills model with CGE (R^2^=0.520 and R^2^=0.529 vs R^2^=0.615). The latter also includes a 5th predictor, MI, but in our models, including MI made very little difference in the value of R^2^. Another notable difference is that the standardized *β* coefficient of the Euclidean distance has the lowest values in our female and male models, while in the Mills model, it has the highest.

Finally, the first principal component of the four vascular features exhibits a Spearman’s rank correlation coefficient of |*ρ*| = 0.449 with the T2:T1 ratio across 38 regions reported in Fulcher et al. (2019), and |*ρ*| = 0.544 with the consensus functional hierarchy gradient. These correlation magnitudes are comparable to those reported for various cortical features in the same study (e.g., interareal axonal connectivity, and densities of glial and inhibitory cells). It should be noted that the first principal component is mostly composed of the tortuosity, the capillary radius and the branching point density.

### 3.3 Whole-brain model of functional connectivity

The functional connectivity models using the 200 whole-brain ROIs and the different combinations of predictors tested are presented in table 3. Across all anesthetized mice models, a decrease in explained variance is observed compared to the models using the 78 cerebrum ROIs. At this scale, the VS explains 10.8% and 9.8% of the variance of the FC of female and male models. These values are still significantly higher than the average Pearson correlation coefficient from the spatial autocorrelation preserving null model ⟨R^2^⟩=0.02, SD=0.01 and ⟨R^2^⟩=0.01, SD=0.01 for female and male datasets (see supp. figure 3 for the correlation values distribution of the null models). Additionally, the addition of vascular similarity to the model still increases the explained variance, albeit its contribution in the full model decreases, as reflected by the values of the standardized coefficients.

**Fig. 3.**
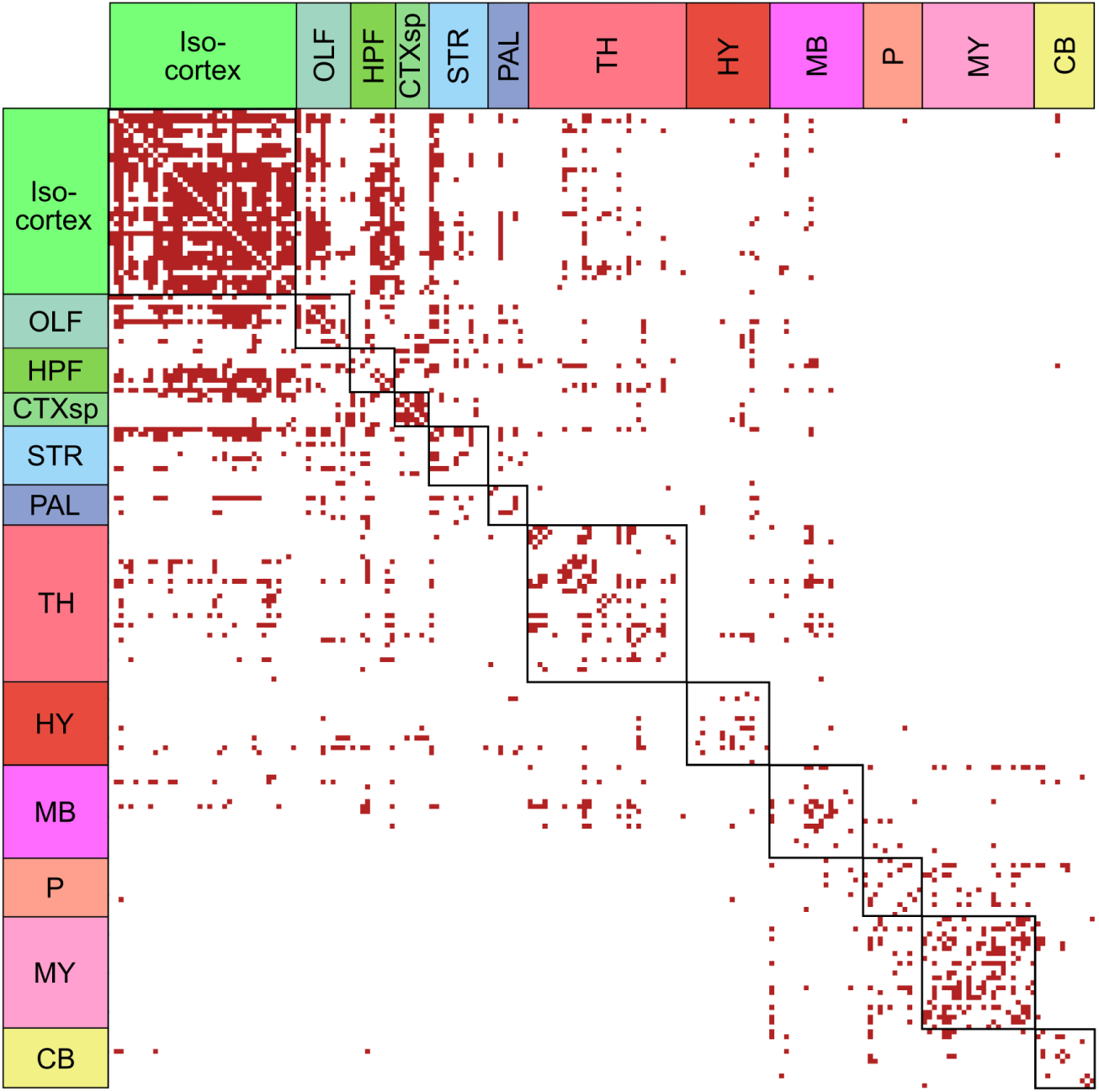
Average binary overlap adjacency matrix between vascular similarity and functional connectivity ordered by the mouse brain 12 major divisions: isocortex, olfactory areas, hippocampus, cortical subplate, striatum, pallidum, thalamus, hypothalamus, midbrain, pons, medulla, and cerebellum. The overlap matrix was made using 10 to 30% connection density. For visualization purposes, the regions in the same major division are outlined by black lines.

**Table 3.** Standardized coefficients *β*, R^2^, and Akaike information criterion (AIC) values for the whole-brain multiple linear models of FC using different predictors. The *β* of each model are in the same order as the model predictors. The AIC is an estimator of prediction error, with lower values indicating a better model.

| Model predictors | Anesthetized females |  |  | Anesthetized males |  |  | Awake males |  |  |
| --- | --- | --- | --- | --- | --- | --- | --- | --- | --- |
| | $\beta$ | $R^2$ | AIC | $\beta$ | $R^2$ | AIC | $\beta$ | $R^2$ | AIC |
| **CO** | 0.512 | 0.262 | 50423 | 0.480 | 0.231 | 51259 | 0.567 | 0.321 | 48276 |
| **VS** | 0.328 | 0.108 | 54213 | 0.313 | 0.098 | 54430 | 0.251 | 0.063 | 54634 |
| **ED** | 0.444 | 0.197 | 52110 | 0.490 | 0.240 | 51023 | 0.589 | 0.347 | 47512 |
| **SA** | 0.475 | 0.225 | 51401 | 0.530 | 0.281 | 49919 | 0.960 | 0.246 | 50359 |
| **ED, SA** | 0.264 | 0.274 | 50102 | 0.286 | 0.338 | 48259 | 0.454 | 0.391 | 46143 |
|  | 0.331 |  |  | 0.374 |  |  | 0.249 |  |  |
| **CO, VS** | 0.461 | 0.308 | 49140 | 0.431 | 0.273 | 50125 | 0.538 | 0.336 | 47838 |
|  | 0.221 |  |  | 0.212 |  |  | 0.126 |  |  |
| **CO, ED, SA** | 0.316 | 0.341 | 48174 | 0.219 | 0.371 | 47265 | 0.318 | 0.459 | 43828 |
|  | 0.162 |  |  | 0.216 |  |  | 0.352 |  |  |
|  | 0.226 |  |  | 0.301 |  |  | 0.144 |  |  |
| **VS, ED, SA** | 0.228 | 0.324 | 48694 | 0.199 | 0.376 | 47088 | 0.127 | 0.406 | 45641 |
|  | 0.239 |  |  | 0.264 |  |  | 0.440 |  |  |
|  | 0.301 |  |  | 0.349 |  |  | 0.233 |  |  |
| **CO, VS, ED, SA** | 0.284 | 0.377 | 47073 | 0.190 | 0.400 | 46318 | 0.302 | 0.467 | 43540 |
|  | 0.195 |  |  | 0.177 |  |  | 0.092 |  |  |
|  | 0.151 |  |  | 0.206 |  |  | 0.092 |  |  |
|  | 0.211 |  |  | 0.288 |  |  | 0.137 |  |  |

To explore the effect of anesthesia, the models were also computed with the awake male dataset. The correlation between the awake and anesthetized male dataset is lower than the correlation between the two anesthetized datasets with r=0.758 and r=0.594 for the cerebrum and the whole-brain respectively. In the whole-brain, VS explains less variance (6.3%) in the awake mouse FC compared to the anesthetized mouse FC. Other notable differences between the anesthetized and awake models are the explained variance by the Euclidean distance (34.7%) and the communicability (32.1%) which have both higher values than in the anesthetized models. The awake model using the four predictors also has higher explained variance (46.7%) than its anesthetized counterparts (R^2^=0.377 for females and R^2^=0.400 for males). The models were also computed for the cerebrum 78 ROIs with results shown in supp. table 1.

Finally, to investigate the decrease of the coefficient of variation in the simple linear regression model of FC using VS in the whole-brain model, an average overlap binary matrix between these two was made using thresholded matrices from 30% to 10% of connection density with 1% interval. The resulting overlap matrix is presented in figure 3. The strongest edges in the VS matrix are mostly the ones connecting two regions in the same major division of the brain. In fact, 32.7% of the top 10% edges with the strongest values are in the same major division. The same phenomenon is observed in functional connectivity where 41.2% of the top 10% strongest edges are in the same major divisions. Partly due to this, 47.9% of the overlap between the functional connectivity and vascular similarity occurs between regions in the same major division despite them constituting only 10.5% of all possible edges. Also, two major divisions that have more than 60% of their edges overlap are the isocortex (65.9%) and the cortical subplate (76.9%) which are all located in the cerebrum.

## 4 Discussion

In this work, the linear relationship between resting-state functional connectivity and vascular similarity was studied in awake and anesthetized C57BL/6J mice. VS is an Euclidean distance-based similarity measure of the vasculature across anatomically defined regions. VS was used in conjunction with commonly used predictors of FC utilizing multiple linear models. This analysis was conducted using FC models composed of two sets of ROIs on female and male resting-state fMRI datasets. Some of the results of this study were compared to those of Mills et al. (2018) in which multiple linear models of FC in mice were also exploited. Finally, the overlap between functional connectivity and vascular similarity was studied.

To begin, single-feature VS was studied. First, comparable R^2^ values were found for both anesthetized FC datasets despite the fact that the whole-brain vasculature dataset was only composed of male subjects. This was not necessarily expected because sex-related differences were found in the mouse brain vasculature (Quintana et al. 2019). Nonetheless, the authors also found that for some regions the similarities and dissimilarities of the vascular profile between subregions were conserved across sexes. Because VS is dependent on the relative difference between the features, the VS matrix could be mostly consistent across sexes. This proximity could also be explained by the strong correlation between the anesthetized male and female FC (r=0.816). It is important to note that all results from this paper were obtained using the C57Bl/6 strain. Significant differences in cerebral vasculature have been documented between mouse strains (Barone et al. 1993; Beckmann 2000), which may limit the generalizability of these findings to other strains.

Upon the results of the single-feature VS models, the four features that would be used in the multiple linear models were the branching point density, the average tortuosity and the average capillary length and radius. The exclusion of vessel and length density from the VS, despite them explaining a similar amount of variance, is coherent because the length density is highly correlated to the BPD, and its use in conjunction with the average capillary length provides a good estimate of the length density. The selected features principally describe the capillary network of the region, including the average tortuosity and the BPD because the capillaries represent the majority of the vessels. In future work, other vascular features could be calculated using graph theory metrics as described in other relevant papers (Blinder et al. 2013; Quintana et al. 2019; Ji et al. 2021; Walek et al. 2023).

Using the 4-features VS in the cerebrum, a significant proportion of the FC variance can be explained in anesthetized females (R^2^=0.159) and males (R^2^=0.151). This proportion diminishes when using all eight features (R^2^=0.121 in females and R^2^=0.110 in males. This could be explained by the addition of features that have significatively lower R^2^ values (i.e. Non-capillary length proportion and proportion of volume filled by vessels). The explained variance of FC using VS alone is smaller than the one computed using simple linear regression with commonly used predictors (communicability, Euclidean distance, spatial adjacency and correlated gene expression). Yet, in the 4-variable models, the substitution of CGE with VS leads to higher explained variance. This is probably due to the collinearity of the variables, the spatial adjacency and the CGE being moderately correlated with the Euclidean distance (r*>*0.6). This makes the Euclidean distance predictor partially redundant in this model, as shown by its standardized coefficient value being at least one order of magnitude lower than other predictors.

When our models are compared to the results of Mills et al. (2018), the variance explained is almost always lower. The only one of our models where explained variance is higher than in the Mills model is the simple linear model using communicability. This difference could be explained by the use of the SC model from Knox et al. (2018). A distinction between their model and the one of Oh et al. (2014) lies in its prediction of minimal yet non-zero connections, in contrast to the latter, which attributes zero weight to such edges. This discrepancy may significantly influence communicability, particularly due to the greater emphasis it places on more direct regional connections. Indeed, the communicability derived from the SC matrix of Oh et al. (2014) explained considerably less variance. However, this could also be due to the fact that the regions composing the Oh SC matrix are defined by the CCFv2, a different version of the atlas than the CCFv3 we used to compute FC (Oh et al. 2014; Wang et al. 2020). In the Mills models, the predictor that contributes the most is the Euclidean distance. In fact, the multivariable model using the Euclidean distance and the spatial adjacency explained more variance than all of our models. This difference could be attributed to the fact that Mills et al. (2018) exclusively used isoflurane, whereas our anesthetized datasets employed a combination of isoflurane and medetomidine. Indeed, it has been shown that the anesthetics can significantly impact the FC (Jonckers et al. 2014; Xie et al. 2020) and that the mixture of isoflurane and medetomidine may have different impacts than using isoflurane alone (Tsurugizawa and Yoshimaru 2021). Furthermore, we observe the same phenomenon in the awake FC which has higher correlation with Euclidean distance in both ROIs subsets compared to our anesthetized dataset. This could be due, in part, to the fact that FC in awake mice shows increased between-network communication (Gutierrez-Barragan et al. 2022). Also, upon a thorough review of the existing literature, we were unable to locate other values of the correlation between the regional Euclidean distance and FC which could allow us to judge the accuracy of these values. Nonetheless, the similar values of the coefficient of determination between the male and female independent anesthetized datasets attest to the reproducibility of this analysis.

The moderately high Spearman correlation between the first principal components of the four selected vascular features and both the T1:T2 ratio (|*ρ*| = 0.449) and the consensus functional hierarchy (|*ρ*| = 0.544) across 38 cortical regions suggests potential alignment between vascular structure and functional hierarchy. This agreement may arise because the vascular architecture could reflect the shifting metabolic demands from the primary sensory regions to the integrative prefrontal areas. As shown in Ji et al. (2021), the mouse brain microvasculature is intrinsically linked to its metabolic load. Furthermore, the vasculature might also act as a marker for the types of cells present in a region, a concept that will be elaborated on later. For certain cell types, this could align with the functional hierarchy (Fulcher et al. 2019).

The addition of VS in the full multivariable linear model using the whole-brain set of 200 ROIs also allows for a more accurate depiction of the anesthetized FC as shown by the R^2^ and AIC values. Nonetheless, we found a decrease in the explained variance of all FC models using this set of ROIs compared to the cerebrum ROIs. This might be accounted for by the fact that almost half of the overlap between FC and VS (47.9%) is between regions in the same major division of the brain. In the subset of 78 regions 28.3% of the possible edges between regions are inside the same major division compared to 10.5% in the full set of 200 regions. Furthermore, the two major brain divisions where the overlap is the highest (the isocortex and the cortical subplate) are also present in the smaller subset. The fact that higher vascular similarity values are observed between regions in the same major division should not come as a surprise. It has been shown that the probability distribution of many features such as vessel length density, branch point density and radius size vary considerably across major regions (Kirst et al. 2020; Ji et al. 2021).

Moreover, we observe that the correlation between FC and VS is weaker in awake mouse, suggesting that anesthetics enhance the FC-VS correlation. This could be attributed to the effect that isoflurane and medetomidine have on vasculature. Indeed, it is known that isoflurane acts as a vasodilatator (Matta et al. 1999) and medetomidine as a vasoconstrictor (Burnside et al. 2013). Due to this, both anesthetics affect the hemodynamics of the brain, notably the cerebral blood flow and cerebrovascular reactivity (Slupe and Kirsch 2018; Munting et al. 2019). These two parameters are known to influence BOLD functional connectivity, as reviewed in Guilbert et al. (2022).

Multiple causes could explain the correlation between the functional connectivity and the vascular variability. Three hypotheses are that the regional vasculature could predict the amplitude of the BOLD signal directly, indirectly or both. In a direct relationship, it is meant that the topology of the local vasculature could influence the amplitude and timing of the BOLD signal measured by fMRI (Buxton 2013; Gagnon et al. 2015; Viessmann et al. 2019). Secondly, the vasculature might serve as an indicator of the types of cells present in a given region. This aspect is significant as neurovascular coupling, and consequently, the amplitude of the BOLD signal is dependent upon the variability of cell types within a region (Drew 2019; Howarth et al. 2020; Tsvetanov et al. 2021). For instance, length density is correlated with pericyte density and inversely correlated with neuronal nitric oxide synthase-positive neurons (nNOS) density (Wu et al. 2022). Both cell types are important in neurovascular coupling, as nNOS impacts blood flow through its vasodilatory effects (Echagarruga et al. 2020; Krawchuk et al. 2020; Lee et al. 2020) and pericytes have a role in the regulation of blood flow (Hall et al. 2014; Alarcon-Martinez et al. 2020). Here, the amplitude of the BOLD signal is related to the measured FC as it influences the Signal-to-Noise Ratio (SNR) and a reduced SNR can diminish the correlation between two time series, potentially leading to lower predictions of FC (Liu 2013; Archila-Meléndez et al. 2020; Guilbert et al. 2022).

Finally, one approach to ascertain whether vasculature variability is intricately linked to neuronal FC or if it primarily affects the magnitude of the BOLD signal, subsequently impacting measured FC, is to explore the correlation between non-BOLD FC and VS. This kind of functional connectivity can be measured using genetically encoded calcium indicators (such as GCaMP) (Tian et al. 2009; Akerboom et al. 2012; Dana et al. 2019) in combination with fluorescence imaging techniques such as widefield imaging. The combination of these techniques has been used in the mouse to image cortical functional connectivity (Silasi et al. 2016; Wright et al. 2017) and even simultaneously with BOLD FC (Lake et al. 2020).

## 5 Conclusion

Vascular similarity seems to explain a significant proportion of the variance of restingstate fMRI-based functional connectivity. More specifically, regions with a similar capillary bed tend to have comparable functional connectivity profiles. Also, the use of VS in multivariable linear models leads to more accurate predictions of FC. This can be attributed partially to the fact that VS is less correlated with Euclidean distance than other commonly used predictors. The overlap between FC and VS seems to be higher within regions of the same major brain division. Finally, according to our results, anesthesia seems to amplify the correlation between VS and FC.

Lastly, in this research, the data utilized was derived from average measurements across multiple subjects. Future studies might benefit from investigating the impact of regional vascular variability on functional connectivity at the individual level. Such an analysis could be facilitated through the acquisition of resting-state fMRI from individual subjects, followed by the application of methodologies described in recent literature to reconstruct the vasculature of the brain (Quintana et al. 2019; Todorov et al. 2020; Kirst et al. 2020; Ji et al. 2021; Wu et al. 2022). This approach could prove valuable in researching neurovascular disorders to study the influence of variables such as sex, age, and disease on an individual’s vascular structure (Quintana et al. 2019; Walek et al. 2023; Meyer et al. 2008) and how it is reflected in whole-brain functional connectivity.

## Acknowledgments

We would like to thank Antoine Légaré, Alexandre Cléroux-Cuillerier, Pierre Girard-Collins and Jérémie Guilbert for their help editing the manuscript.

## Statements and Declarations

### Author Contributions

All authors contributed to the study conception and design. Data analysis was performed by François Gaudreault. The first draft of the manuscript was written by François Gaudreault and all authors commented on previous versions of the manuscript. All authors read and approved the final manuscript.

### Data Availability

All the raw datasets used in this study are openly available or can be accessed upon request to the authors. The processed data and source code used in this study are available from the corresponding author on reasonable request.

## Footnotes

1 https://openneuro.org/datasets/ds001653/versions/1.0.2 aged between 10 and 15 weeks and two datasets of adult (< 6 months) male C57BL/6J mice from Gutierrez-Barragan, et al. (2022), one with N = 14 anesthetized males and another of N=10 awake males. Most analyses in this paper focus on the anesthetized datasets; when referring to the unanesthetized dataset, we will call it the awake dataset.

2 https://github.com/CoBrALab/RABIES

3 https://github.com/CoBrALab/optimized_antsMultivariateTemplateConstruction

4 http://www.brain-map.org/

5 https://connectivity.brain-map.org/

## References

Akerboom, J., T.W. Chen, T.J. Wardill, L. Tian, J.S. Marvin, S. Mutlu, N.C. Calderón, F. Esposti, B.G. Borghuis, X.R. Sun, A. Gordus, M.B. Orger, R. Portugues, F. Engert, J.J. Macklin, A. Filosa, A. Aggarwal, R.A. Kerr, R. Takagi, S. Kracun, E. Shigetomi, B.S. Khakh, H. Baier, L. Lagnado, S.S.H. Wang, C.I. Bargmann, B.E. Kimmel, V. Jayaraman, K. Svoboda, D.S. Kim, E.R. Schreiter, and L.L. Looger. 2012, October. Optimization of a GCaMP Calcium Indicator for Neural Activity Imaging. Journal of Neuroscience 32 (40): 13819–13840. 10.1523/JNEUROSCI.2601-12.2012.

Alarcon-Martinez, L., D. Villafranca-Baughman, H. Quintero, J.B. Kacerovsky, F. Dotigny, K.K. Murai, A. Prat, P. Drapeau, and A. Di Polo. 2020, September. Interpericyte tunnelling nanotubes regulate neurovascular coupling. Nature 585 (7823): 91–95. 10.1038/s41586-020-2589-x.

Archila-Meléndez, M.E., C. Sorg, and C. Preibisch. 2020, September. Modeling the impact of neurovascular coupling impairments on BOLD-based functional connectivity at rest. NeuroImage 218: 116871. 10.1016/j.neuroimage.2020.116871.

Attwell, D. and C. Iadecola. 2002, December. The neural basis of functional brain imaging signals. Trends in Neurosciences 25 (12): 621–625. 10.1016/S0166-2236(02)02264-6.

Avants, B.B., N.J. Tustison, G. Song, P.A. Cook, A. Klein, and J.C. Gee. 2011, February. A reproducible evaluation of ANTs similarity metric performance in brain image registration. NeuroImage 54 (3): 2033–2044. 10.1016/j.neuroimage.2010.09.025.

Bandettini, P.A., E.C. Wong, R.S. Hinks, R.S. Tikofsky, and J.S. Hyde. 1992. Time course EPI of human brain function during task activation. Magnetic Resonance in Medicine 25 (2): 390–397. 10.1002/mrm.1910250220.

Barone, F.C., D.J. Knudsen, A.H. Nelson, G.Z. Feuerstein, and R.N. Willette. 1993, July. Mouse Strain Differences in Susceptibility to Cerebral Ischemia are Related to Cerebral Vascular Anatomy. Journal of Cerebral Blood Flow & Metabolism 13 (4): 683–692. 10.1038/jcbfm.1993.87.

Beckmann, N. 2000. High resolution magnetic resonance angiography non-invasively reveals mouse strain differences in the cerebrovascular anatomy in vivo. Magnetic Resonance in Medicine 44 (2): 252–258. 10.1002/1522-2594(200008)44:2⟨252::AID-MRM12⟩3.0.CO;2-G.

Belliveau, J.W., D.N. Kennedy, R.C. McKinstry, B.R. Buchbinder, R.M. Weisskoff, M.S. Cohen, J.M. Vevea, T.J. Brady, and B.R. Rosen. 1991, November. Functional mapping of the human visual cortex by magnetic resonance imaging. *Science (New York*, N.Y.) 254 (5032): 716–719. 10.1126/science.1948051.

Blinder, P., P.S. Tsai, J.P. Kaufhold, P.M. Knutsen, H. Suhl, and D. Kleinfeld. 2013, July. The cortical angiome: an interconnected vascular network with noncolumnar patterns of blood flow. Nature Neuroscience 16 (7): 889–897. 10.1038/nn.3426.

Burnside, W.M., P.A. Flecknell, A.I. Cameron, and A.A. Thomas. 2013, December. A comparison of medetomidine and its active enantiomer dexmedetomidine when administered with ketamine in mice. BMC Veterinary Research 9 (1): 48. 10.1186/1746-6148-9-48.

Burt, J.B., M. Helmer, M. Shinn, A. Anticevic, and J.D. Murray. 2020, October. Generative modeling of brain maps with spatial autocorrelation. NeuroImage 220: 117038. 10.1016/j.neuroimage.2020.117038.

Buxton, R.B. 2013, September. The physics of functional magnetic resonance imaging (fMRI). Reports on Progress in Physics 76 (9): 096601. 10.1088/0034-4885/76/9/096601.

Chen, Z., A. Caprihan, and V. Calhoun. 2010. Effect of surrounding vasculature on intravoxel BOLD signal. Medical Physics 37 (4): 1778–1787. 10.1118/1.3366251.

Coletta, L., M. Pagani, J.D. Whitesell, J.A. Harris, B. Bernhardt, and A. Gozzi. 2020, December. Network structure of the mouse brain connectome with voxel resolution. Science Advances 6 (51): eabb7187. 10.1126/sciadv.abb7187.

Cox, R.W. 1996, June. AFNI: software for analysis and visualization of functional magnetic resonance neuroimages. Computers and Biomedical Research, an International Journal 29 (3): 162–173. 10.1006/cbmr.1996.0014.

Crofts, J.J. and D.J. Higham. 2009, April. A weighted communicability measure applied to complex brain networks. Journal of the Royal Society Interface 6 (33): 411–414. 10.1098/rsif.2008.0484.

Damoiseaux, J.S. and M.D. Greicius. 2009, October. Greater than the sum of its parts: a review of studies combining structural connectivity and resting-state functional connectivity. Brain Structure and Function 213 (6): 525–533. 10.1007/s00429-009-0208-6.

Dana, H., Y. Sun, B. Mohar, B.K. Hulse, A.M. Kerlin, J.P. Hasseman, G. Tsegaye, A. Tsang, A. Wong, R. Patel, J.J. Macklin, Y. Chen, A. Konnerth, V. Jayaraman, L.L. Looger, E.R. Schreiter, K. Svoboda, and D.S. Kim. 2019, July. High-performance calcium sensors for imaging activity in neuronal populations and microcompartments. Nature Methods 16 (7): 649–657. 10.1038/s41592-019-0435-6.

Desrosiers-Grégoire, G., G.A. Devenyi, J. Grandjean, and M.M. Chakravarty. 2024, August. A standardized image processing and data quality platform for rodent fMRI. Nature Communications 15 (1): 6708. 10.1038/s41467-024-50826-8.

Dorr, A.E., J.P. Lerch, S. Spring, N. Kabani, and R.M. Henkelman. 2008, August. High resolution three-dimensional brain atlas using an average magnetic resonance image of 40 adult C57Bl/6J mice. NeuroImage 42 (1): 60–69. 10.1016/j.neuroimage.2008.03.037.

Drew, P.J. 2019, October. Vascular and neural basis of the BOLD signal. Current Opinion in Neurobiology 58: 61–69. 10.1016/j.conb.2019.06.004.

Echagarruga, C.T., K.W. Gheres, J.N. Norwood, and P.J. Drew. 2020, October. nNOS-expressing interneurons control basal and behaviorally evoked arterial dilation in somatosensory cortex of mice. eLife 9: e60533. 10.7554/eLife.60533.

Fulcher, B.D., J.D. Murray, V. Zerbi, and X.J. Wang. 2019, March. Multimodal gradients across mouse cortex. Proceedings of the National Academy of Sciences of the United States of America 116 (10): 4689–4695. 10.1073/pnas.1814144116.

Gagnon, L., S. Sakadžić, F. Lesage, J.J. Musacchia, J. Lefebvre, Q. Fang, M.A. Yücel, K.C. Evans, E.T. Mandeville, J. Cohen-Adad, J.R. Polimeni, M.A. Yaseen, E.H. Lo, D.N. Greve, R.B. Buxton, A.M. Dale, A. Devor, and D.A. Boas. 2015, February. Quantifying the Microvascular Origin of BOLD-fMRI from First Principles with Two-Photon Microscopy and an Oxygen-Sensitive Nanoprobe. Journal of Neuroscience 35 (8): 3663–3675. 10.1523/JNEUROSCI.3555-14.2015.

Goñi, J., M.P. van den Heuvel, A. Avena-Koenigsberger, N. Velez de Mendizabal, R.F. Betzel, A. Griffa, P. Hagmann, B. Corominas-Murtra, J.P. Thiran, and O. Sporns. 2014, January. Resting-brain functional connectivity predicted by analytic measures of network communication. Proceedings of the National Academy of Sciences 111 (2): 833–838. 10.1073/pnas.1315529111.

Grandjean, J., V. Zerbi, J.H. Balsters, N. Wenderoth, and M. Rudin. 2017, August. Structural Basis of Large-Scale Functional Connectivity in the Mouse. Journal of Neuroscience 37 (34): 8092–8101. 10.1523/JNEUROSCI.0438-17.2017.

Greicius, M.D., K. Supekar, V. Menon, and R.F. Dougherty. 2009, January. Resting-State Functional Connectivity Reflects Structural Connectivity in the Default Mode Network. Cerebral Cortex 19 (1): 72–78. 10.1093/cercor/bhn059.

Guilbert, J., A. Légaré, P.D. Koninck, P. Desrosiers, and M. Desjardins. 2022, April. Toward an integrative neurovascular framework for studying brain networks. Neurophotonics 9 (3): 032211. 10.1117/1.NPh.9.3.032211.

Gutierrez-Barragan, D., N.A. Singh, F.G. Alvino, L. Coletta, F. Rocchi, E. De Guzman, A. Galbusera, M. Uboldi, S. Panzeri, and A. Gozzi. 2022, February. Unique spatiotemporal fMRI dynamics in the awake mouse brain. Current Biology 32 (3): 631–644.e6. 10.1016/j.cub.2021.12.015.

Hall, C.N., C. Reynell, B. Gesslein, N.B. Hamilton, A. Mishra, B.A. Sutherland, F.M. O’Farrell, A.M. Buchan, M. Lauritzen, and D. Attwell. 2014, April. Capillary pericytes regulate cerebral blood flow in health and disease. Nature 508 (7494): 55–60. 10.1038/nature13165.

Honey, C.J., O. Sporns, L. Cammoun, X. Gigandet, J.P. Thiran, R. Meuli, and P. Hagmann. 2009, February. Predicting human resting-state functional connectivity from structural connectivity. Proceedings of the National Academy of Sciences 106 (6): 2035–2040. 10.1073/pnas.0811168106.

Howarth, C., A. Mishra, and C.N. Hall. 2020, November. More than just summed neuronal activity: how multiple cell types shape the BOLD response. Philosophical Transactions of the Royal Society B: Biological Sciences 376 (1815): 20190630. 10.1098/rstb.2019.0630.

Howe, F.A., A.G. Filler, B.A. Bell, and J.R. Griffiths. 1992. Magnetic Resonance Neurography. Magnetic Resonance in Medicine 28 (2): 328–338. 10.1002/mrm.1910280215.

Huntenburg, J.M., P.L. Bazin, and D.S. Margulies. 2018, January. Large-Scale Gradients in Human Cortical Organization. Trends in Cognitive Sciences 22 (1): 21–31. 10.1016/j.tics.2017.11.002.

Ji, X., T. Ferreira, B. Friedman, R. Liu, H. Liechty, E. Bas, J. Chandrashekar, and D. Kleinfeld. 2021, April. Brain microvasculature has a common topology with local differences in geometry that match metabolic load. Neuron 109 (7): 1168–1187.e13. 10.1016/j.neuron.2021.02.006.

Jonckers, E., R. Delgado y Palacios, D. Shah, C. Guglielmetti, M. Verhoye, and A. Van der Linden. 2014, October. Different anesthesia regimes modulate the functional connectivity outcome in mice. Magnetic Resonance in Medicine 72 (4): 1103–1112. 10.1002/mrm.24990.

Kirst, C., S. Skriabine, A. Vieites-Prado, T. Topilko, P. Bertin, G. Gerschenfeld, F. Verny, P. Topilko, N. Michalski, M. Tessier-Lavigne, and N. Renier. 2020, February. Mapping the Fine-Scale Organization and Plasticity of the Brain Vasculature. Cell 180 (4): 780–795.e25. 10.1016/j.cell.2020.01.028.

Knox, J.E., K.D. Harris, N. Graddis, J.D. Whitesell, H. Zeng, J.A. Harris, E. Shea-Brown, and S. Mihalas. 2018, December. High-resolution data-driven model of the mouse connectome. Network Neuroscience 3 (1): 217–236. 10.1162/netn_a_00066.

Koch, M.A., D.G. Norris, and M. Hund-Georgiadis. 2002, May. An Investigation of Functional and Anatomical Connectivity Using Magnetic Resonance Imaging. NeuroImage 16 (1): 241–250. 10.1006/nimg.2001.1052.

Krawchuk, M.B., C.F. Ruff, X. Yang, S.E. Ross, and A.L. Vazquez. 2020, July. Optogenetic assessment of VIP, PV, SOM and NOS inhibitory neuron activity and cerebral blood flow regulation in mouse somato-sensory cortex. Journal of Cerebral Blood Flow & Metabolism 40 (7): 1427–1440. 10.1177/0271678X19870105.

Kwong, K.K., J.W. Belliveau, D.A. Chesler, I.E. Goldberg, R.M. Weisskoff, B.P. Poncelet, D.N. Kennedy, B.E. Hoppel, M.S. Cohen, and R. Turner. 1992, June. Dynamic magnetic resonance imaging of human brain activity during primary sensory stimulation. Proceedings of the National Academy of Sciences 89 (12): 5675–5679. 10.1073/pnas.89.12.5675.

Lake, E.M.R., X. Ge, X. Shen, P. Herman, F. Hyder, J.A. Cardin, M.J. Higley, D. Scheinost, X. Papademetris, M.C. Crair, and R.T. Constable. 2020, December. Simultaneous cortex-wide fluorescence Ca2+ imaging and whole-brain fMRI. Nature Methods 17 (12): 1262–1271. 10.1038/s41592-020-00984-6.

Lee, L., L. Boorman, E. Glendenning, C. Christmas, P. Sharp, P. Redgrave, O. Shabir, E. Bracci, J. Berwick, and C. Howarth. 2020, April. Key Aspects of Neurovascular Control Mediated by Specific Populations of Inhibitory Cortical Interneurons. Cerebral Cortex 30 (4): 2452–2464. 10.1093/cercor/bhz251.

Lein, E.S., M.J. Hawrylycz, N. Ao, M. Ayres, A. Bensinger, A. Bernard, A.F. Boe, M.S. Boguski, K.S. Brockway, E.J. Byrnes, L. Chen, L. Chen, T.M. Chen, M. Chi Chin, J. Chong, B.E. Crook, A. Czaplinska, C.N. Dang, S. Datta, N.R. Dee, A.L. Desaki, T. Desta, E. Diep, T.A. Dolbeare, M.J. Donelan, H.W. Dong, J.G. Dougherty, B.J. Duncan, A.J. Ebbert, G. Eichele, L.K. Estin, C. Faber, B.A. Facer, R. Fields, S.R. Fischer, T.P. Fliss, C. Frensley, S.N. Gates, K.J. Glattfelder, K.R. Halverson, M.R. Hart, J.G. Hohmann, M.P. Howell, D.P. Jeung, R.A. Johnson, P.T. Karr, R. Kawal, J.M. Kidney, R.H. Knapik, C.L. Kuan, J.H. Lake, A.R. Laramee, K.D. Larsen, C. Lau, T.A. Lemon, A.J. Liang, Y. Liu, L.T. Luong, J. Michaels, J.J. Morgan, R.J. Morgan, M.T. Mortrud, N.F. Mosqueda, L.L. Ng, R. Ng, G.J. Orta, C.C. Overly, T.H. Pak, S.E. Parry, S.D. Pathak, O.C. Pearson, R.B. Puchalski, Z.L. Riley, H.R. Rockett, S.A. Rowland, J.J. Royall, M.J. Ruiz, N.R. Sarno, K. Schaffnit, N.V. Shapovalova, T. Sivisay, C.R. Slaughterbeck, S.C. Smith, K.A. Smith, B.I. Smith, A.J. Sodt, N.N. Stewart, K.R. Stumpf, S.M. Sunkin, M. Sutram, A. Tam, C.D. Teemer, C. Thaller, C.L. Thompson, L.R. Varnam, A. Visel, R.M. Whitlock, P.E. Wohnoutka, C.K. Wolkey, V.Y. Wong, M. Wood, M.B. Yaylaoglu, R.C. Young, B.L. Youngstrom, X. Feng Yuan, B. Zhang, T.A. Zwingman, and A.R. Jones. 2007, January. Genome-wide atlas of gene expression in the adult mouse brain. Nature 445 (7124): 168–176. 10.1038/nature05453.

Liu, T.T. 2013, October. Neurovascular factors in resting-state functional MRI. NeuroImage 80: 339–348. 10.1016/j.neuroimage.2013.04.071.

Mahani, F.S.N., A. Kalantari, G.R. Fink, M. Hoehn, and M. Aswendt. 2023. A systematic review of the relationship between magnetic resonance imaging based resting-state and structural networks in the rodent brain. Frontiers in Neuroscience 17.

Matta, B.F., K.J. Heath, K. Tipping, and A.C. Summors. 1999, September. Direct cerebral vasodilatory effects of sevoflurane and isoflurane. Anesthesiology 91 (3): 677–680. 10.1097/00000542-199909000-00019.

Mesulam, M.M. 1998, June. From sensation to cognition. Brain: A Journal of Neurology 121 ( Pt 6): 1013–1052. 10.1093/brain/121.6.1013.

Meyer, E.P., A. Ulmann-Schuler, M. Staufenbiel, and T. Krucker. 2008, March. Altered morphology and 3D architecture of brain vasculature in a mouse model for Alzheimer’s disease. Proceedings of the National Academy of Sciences 105 (9): 3587–3592. 10.1073/pnas.0709788105.

Mills, B.D., D.S. Grayson, A. Shunmugavel, O. Miranda-Dominguez, E. Feczko, E. Earl, K.A. Neve, and D.A. Fair. 2018, June. Correlated Gene Expression and Anatomical Communication Support Synchronized Brain Activity in the Mouse Functional Connectome. Journal of Neuroscience 38 (25): 5774–5787. 10.1523/JNEUROSCI.2910-17.2018.

Munting, L.P., M.P. Derieppe, E. Suidgeest, B. Denis de Senneville, J.A. Wells, and L. van der Weerd. 2019. Influence of different isoflurane anesthesia protocols on murine cerebral hemodynamics measured with pseudo-continuous arterial spin labeling. NMR in Biomedicine 32 (8): e4105. 10.1002/nbm.4105.

Ogawa, S., T.M. Lee, A.R. Kay, and D.W. Tank. 1990, December. Brain magnetic resonance imaging with contrast dependent on blood oxygenation. Proceedings of the National Academy of Sciences 87 (24): 9868–9872. 10.1073/pnas.87.24.9868.

Ogawa, S., D.W. Tank, R. Menon, J.M. Ellermann, S.G. Kim, H. Merkle, and K. Ugurbil. 1992, July. Intrinsic signal changes accompanying sensory stimulation: functional brain mapping with magnetic resonance imaging. Proceedings of the National Academy of Sciences 89 (13): 5951–5955. 10.1073/pnas.89.13.5951.

Oh, S.W., J.A. Harris, L. Ng, B. Winslow, N. Cain, S. Mihalas, Q. Wang, C. Lau, L. Kuan, A.M. Henry, M.T. Mortrud, B. Ouellette, T.N. Nguyen, S.A. Sorensen, C.R. Slaughterbeck, W. Wakeman, Y. Li, D. Feng, A. Ho, E. Nicholas, K.E. Hirokawa, P. Bohn, K.M. Joines, H. Peng, M.J. Hawrylycz, J.W. Phillips, J.G. Hohmann, P. Wohnoutka, C.R. Gerfen, C. Koch, A. Bernard, C. Dang, A.R. Jones, and H. Zeng. 2014, April. A mesoscale connectome of the mouse brain. Nature 508 (7495): 207–214. 10.1038/nature13186.

Pierpaoli, C., P. Jezzard, P.J. Basser, A. Barnett, and G. Di Chiro. 1996, December. Diffusion tensor MR imaging of the human brain. Radiology 201 (3): 637–648. 10.1148/radiology.201.3.8939209.

Power, J.D., K.A. Barnes, A.Z. Snyder, B.L. Schlaggar, and S.E. Petersen. 2012, February. Spurious but systematic correlations in functional connectivity MRI networks arise from subject motion. Neuroimage 59 (3): 2142–2154. 10.1016/j.neuroimage.2011.10.018.

Pruim, R.H.R., M. Mennes, D. van Rooij, A. Llera, J.K. Buitelaar, and C.F. Beckmann. 2015, May. ICA-AROMA: A robust ICA-based strategy for removing motion artifacts from fMRI data. NeuroImage 112: 267–277. 10.1016/j.neuroimage.2015.02.064.

Quintana, D.D., S.E. Lewis, Y. Anantula, J.A. Garcia, S.N. Sarkar, J.Z. Cavendish, C.M. Brown, and J.W. Simpkins. 2019, November. The cerebral angiome: High resolution MicroCT imaging of the whole brain cerebrovasculature in female and male mice. NeuroImage 202: 116109. 10.1016/j.neuroimage.2019.116109.

Richards, K., C. Watson, R.F. Buckley, N.D. Kurniawan, Z. Yang, M.D. Keller, R. Beare, P.F. Bartlett, G.F. Egan, G.J. Galloway, G. Paxinos, S. Petrou, and D.C. Reutens. 2011, October. Segmentation of the mouse hippocampal formation in magnetic resonance images. NeuroImage 58 (3): 732–740. 10.1016/j.neuroimage.2011.06.025.

Silasi, G., D. Xiao, M.P. Vanni, A.C.N. Chen, and T.H. Murphy. 2016, July. Intact skull chronic windows for mesoscopic wide-field imaging in awake mice. Journal of Neuroscience Methods 267: 141–149. 10.1016/j.jneumeth.2016.04.012.

Slupe, A.M. and J.R. Kirsch. 2018, December. Effects of anesthesia on cerebral blood flow, metabolism, and neuroprotection. Journal of Cerebral Blood Flow & Metabolism 38 (12): 2192–2208. 10.1177/0271678X18789273.

Steadman, P.E., J. Ellegood, K.U. Szulc, D.H. Turnbull, A.L. Joyner, R.M. Henkelman, and J.P. Lerch. 2014. Genetic Effects on Cerebellar Structure Across Mouse Models of Autism Using a Magnetic Resonance Imaging Atlas. Autism Research 7 (1): 124–137. 10.1002/aur.1344.

Tian, L., S.A. Hires, T. Mao, D. Huber, M.E. Chiappe, S.H. Chalasani, L. Petreanu, J. Akerboom, S.A. McKinney, E.R. Schreiter, C.I. Bargmann, V. Jayaraman, K. Svoboda, and L.L. Looger. 2009, December. Imaging neural activity in worms, flies and mice with improved GCaMP calcium indicators. Nature Methods 6 (12): 875–881. 10.1038/nmeth.1398.

Todorov, M.I., J.C. Paetzold, O. Schoppe, G. Tetteh, S. Shit, V. Efremov, K. Todorov-Völgyi, M. Düring, M. Dichgans, M. Piraud, B. Menze, and A. Ertürk. 2020, April. Machine learning analysis of whole mouse brain vasculature. Nature Methods 17 (4): 442–449. 10.1038/s41592-020-0792-1.

Tsurugizawa, T. and D. Yoshimaru. 2021, November. Impact of anesthesia on static and dynamic functional connectivity in mice. NeuroImage 241: 118413. 10.1016/j.neuroimage.2021.118413.

Tsvetanov, K.A., R.N.A. Henson, and J.B. Rowe. 2021, January. Separating vascular and neuronal effects of age on fMRI BOLD signals. Philosophical Transactions of the Royal Society B: Biological Sciences 376 (1815): 20190631. 10.1098/rstb.2019.0631.

Ullmann, J.F.P., C. Watson, A.L. Janke, N.D. Kurniawan, and D.C. Reutens. 2013, September. A segmentation protocol and MRI atlas of the C57BL/6J mouse neocortex. NeuroImage 78: 196–203. 10.1016/j.neuroimage.2013.04.008.

Viessmann, O., K. Scheffler, M. Bianciardi, L.L. Wald, and J.R. Polimeni. 2019, August. Dependence of resting-state fMRI fluctuation amplitudes on cerebral cortical orientation relative to the direction of B0 and anatomical axes. NeuroImage 196: 337–350. 10.1016/j.neuroimage.2019.04.036.

Vigneau-Roy, N., M. Bernier, M. Descoteaux, and K. Whittingstall. 2014. Regional variations in vascular density correlate with resting-state and task-evoked blood oxygen level-dependent signal amplitude. Human Brain Mapping 35 (5): 1906–1920. 10.1002/hbm.22301.

Vázquez-Rodríguez, B., L.E. Suárez, R.D. Markello, G. Shafiei, C. Paquola, P. Hagmann, M.P. van den Heuvel, B.C. Bernhardt, R.N. Spreng, and B. Misic. 2019, October. Gradients of structure–function tethering across neocortex. Proceedings of the National Academy of Sciences 116 (42): 21219–21227. 10.1073/pnas.1903403116.

Walek, K.W., S. Stefan, J.H. Lee, P. Puttigampala, A.H. Kim, S.W. Park, P.J. Marchand, F. Lesage, T. Liu, Y.W.A. Huang, D.A. Boas, C. Moore, and J. Lee. 2023, May. Near-lifespan longitudinal tracking of brain microvascular morphology, topology, and flow in male mice. Nature Communications 14 (1): 2982. 10.1038/s41467-023-38609-z.

Wang, Q., S.L. Ding, Y. Li, J. Royall, D. Feng, P. Lesnar, N. Graddis, M. Naeemi, B. Facer, A. Ho, T. Dolbeare, B. Blanchard, N. Dee, W. Wakeman, K.E. Hirokawa, A. Szafer, S.M. Sunkin, S.W. Oh, A. Bernard, J.W. Phillips, M. Hawrylycz, C. Koch, H. Zeng, J.A. Harris, and L. Ng. 2020, May. The Allen Mouse Brain Common Coordinate Framework: A 3D Reference Atlas. Cell 181 (4): 936–953.e20. 10.1016/j.cell.2020.04.007.

Wang, S., D.J. Peterson, J.C. Gatenby, W. Li, T.J. Grabowski, and T.M. Madhyastha. 2017. Evaluation of Field Map and Nonlinear Registration Methods for Correction of Susceptibility Artifacts in Diffusion MRI. Frontiers in Neuroinformatics 11: 17. 10.3389/fninf.2017.00017.

Wright, P.W., L.M. Brier, A.Q. Bauer, G.A. Baxter, A.W. Kraft, M.D. Reisman, A.R. Bice, A.Z. Snyder, J.M. Lee, and J.P. Culver. 2017, October. Functional connectivity structure of cortical calcium dynamics in anesthetized and awake mice. PLOS ONE 12 (10): e0185759. 10.1371/journal.pone.0185759.

Wu, Y.t., H.C. Bennett, U. Chon, D.J. Vanselow, Q. Zhang, R. Muñoz-Castañeda, K.C. Cheng, P. Osten, P.J. Drew, and Y. Kim. 2022, June. Quantitative relationship between cerebrovascular network and neuronal cell types in mice. Cell reports 39 (12): 110978. 10.1016/j.celrep.2022.110978.

Xie, H., D.Y. Chung, S. Kura, K. Sugimoto, S.A. Aykan, Y. Wu, S. Sakadžić, M.A. Yaseen, D.A. Boas, and C. Ayata. 2020, April. Differential effects of anesthetics on resting state functional connectivity in the mouse. Journal of Cerebral Blood Flow & Metabolism 40 (4): 875–884. 10.1177/0271678X19847123.

Zamani Esfahlani, F., J. Faskowitz, J. Slack, B. Mišić, and R.F. Betzel. 2022, April. Local structure-function relationships in human brain networks across the lifespan. Nature Communications 13 (1): 2053. 10.1038/s41467-022-29770-y.

