## Supplementary materials for "Microvascular structure variability explains variance in fMRI functional connectivity"

### 1 Supplementary figures and tables

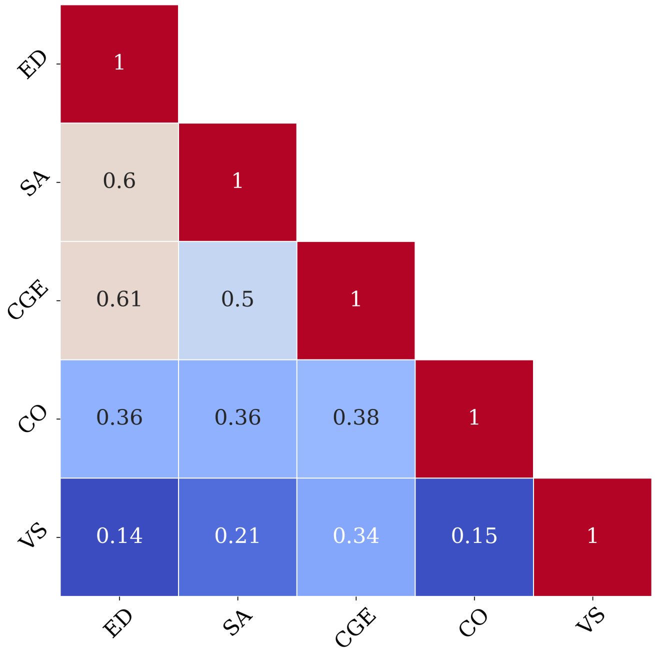


Figure 1 – Correlation matrix between the different predictors of the cerebrum model. ED is the inverse of the spatial Euclidean distance, SA is the spatial adjacency, CGE is the correlated gene expressions, CO is the communicability, and VS is the vascular similarity.

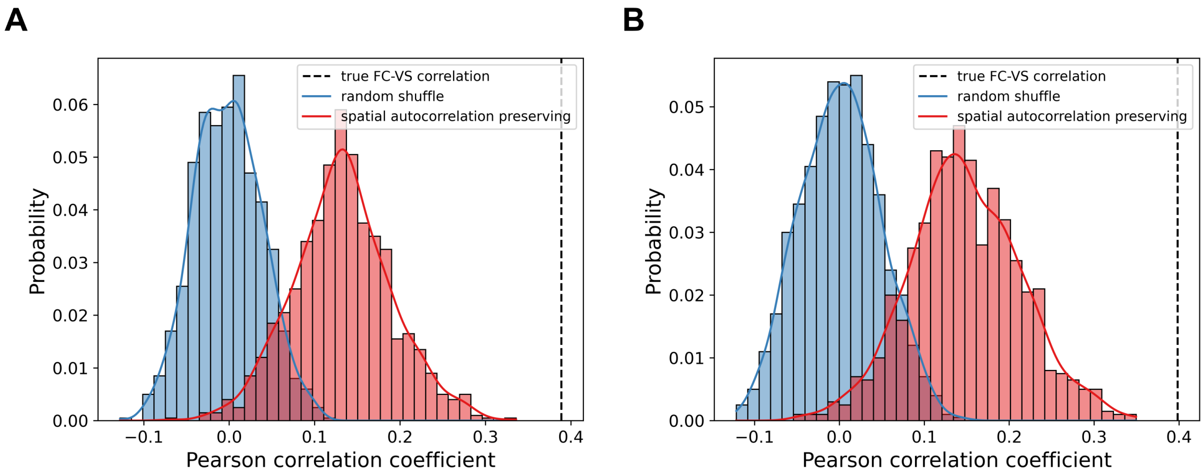


FIGURE 2 – Comparison between the distribution of the correlation values of the random shuffle, the spatial autocorrelation preserving null model distribution and the true correlation between functional connectivity and vascular similarity in the cerebrum. A) The correlation between group anesthetized male FC and VS. B) The correlation between group anesthetized female FC and VS.

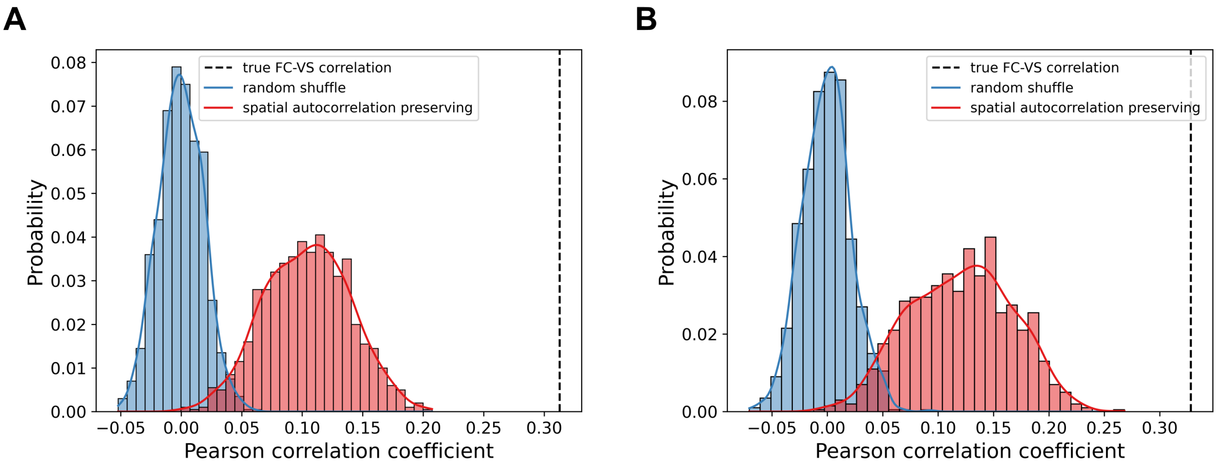


FIGURE 3 – Comparison between the distribution of the correlation values of the random shuffle, the spatial autocorrelation preserving null model distribution and the true correlation between functional connectivity and vascular similarity in the whole brain. A) The correlation between group anesthetized male FC and VS. B) The correlation between group anesthetized female FC and VS.

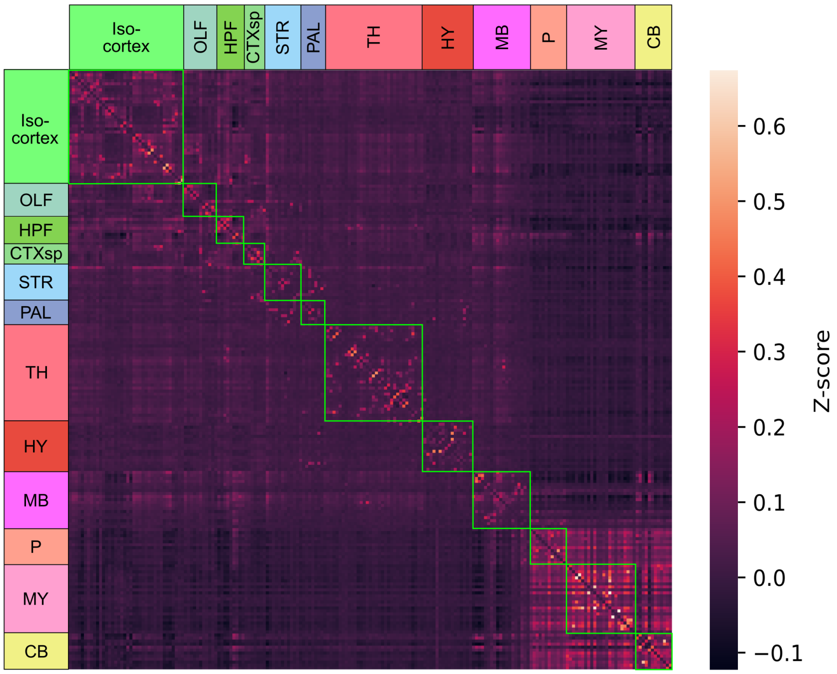


FIGURE 4 – Whole-brain awake male functional connectivity matrix, the ROIs are ordered as in section 2.

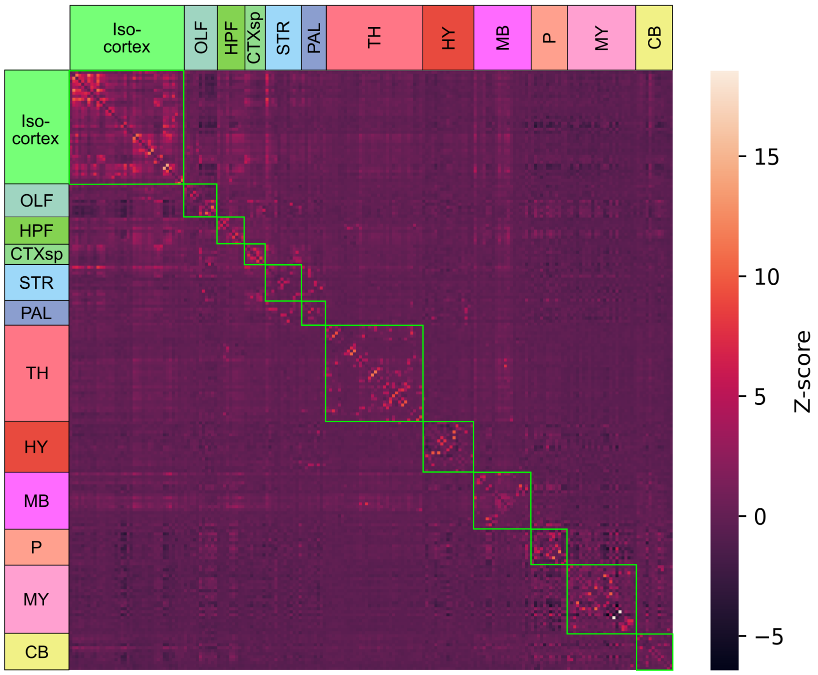


FIGURE 5 – Whole-brain anesthetized male functional connectivity matrix, the ROIs are ordered as in section 2.

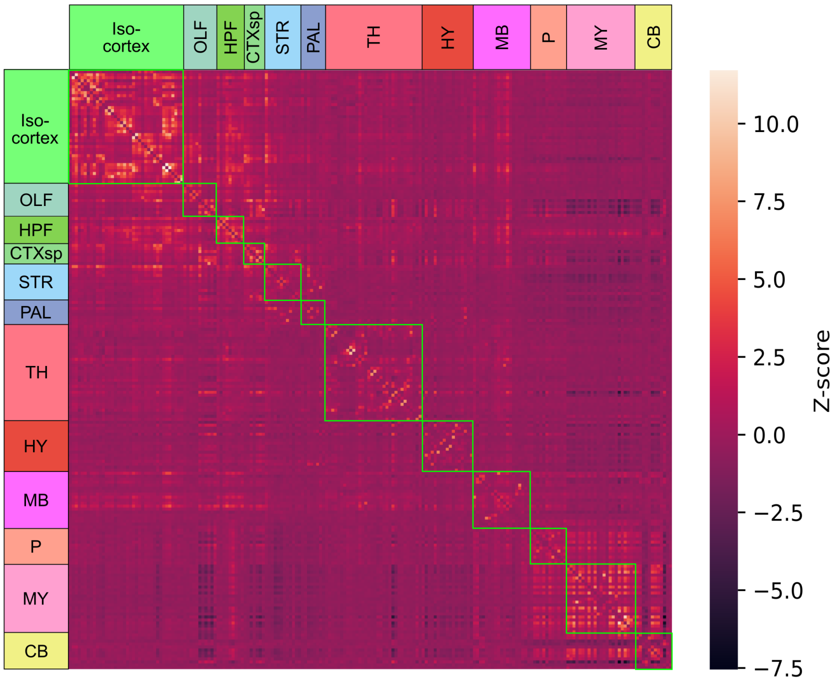


FIGURE 6 – Whole-brain anesthetized female functional connectivity matrix, the ROIs are ordered as in section 2.

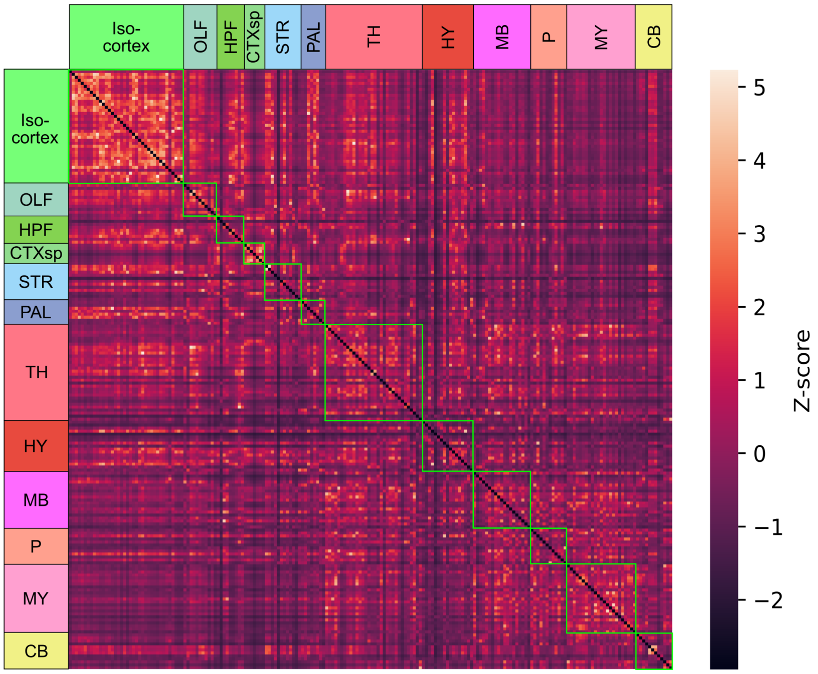


FIGURE 7 – Whole-brain vascular similarity matrix, the ROIs are ordered as in section 2.

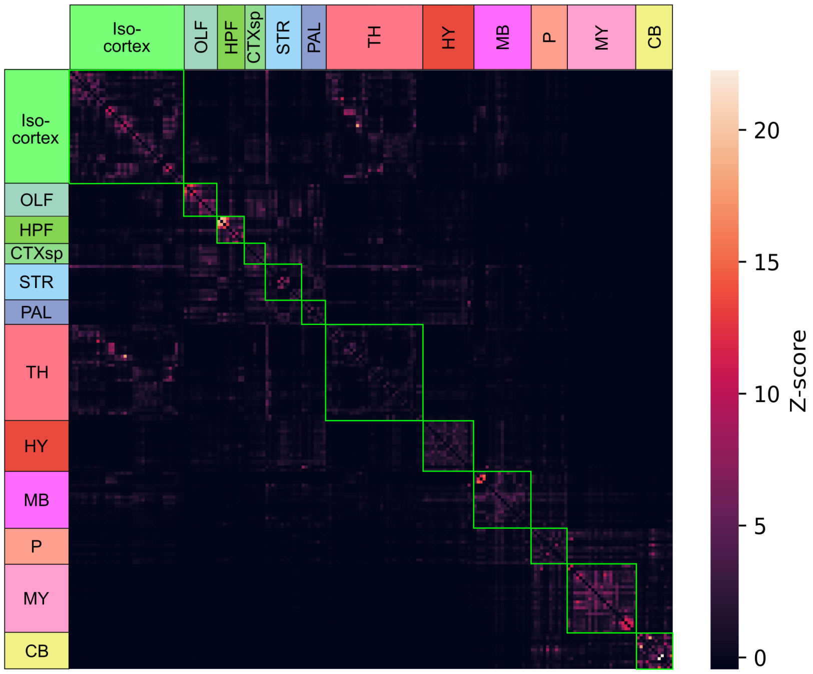


FIGURE 8 – Whole-brain communicability matrix, the ROIs are ordered as in section 2.

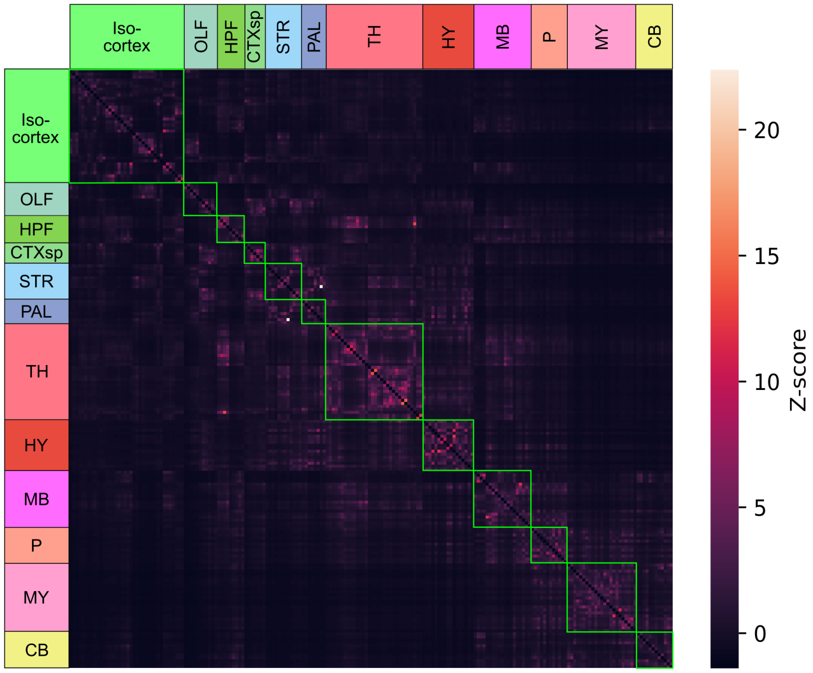


FIGURE 9 – Whole-brain Euclidean distance matrix, the ROIs are ordered as in section 2.

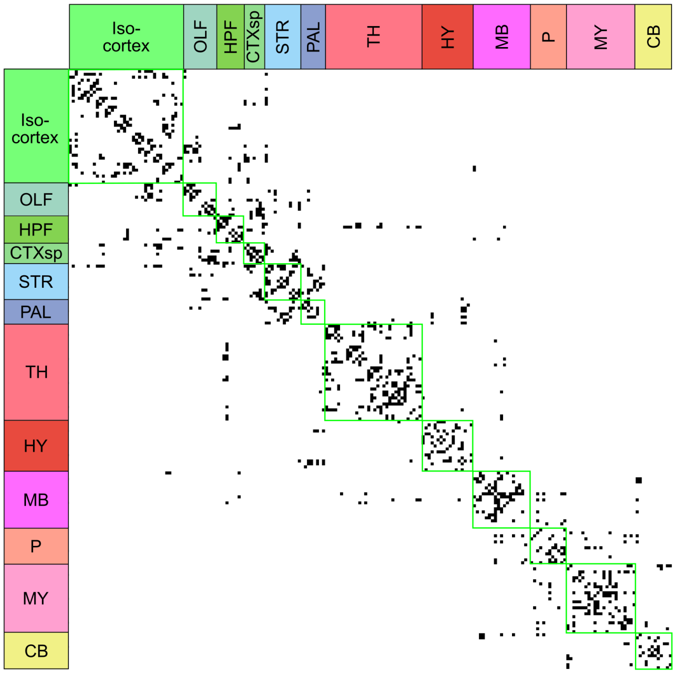


FIGURE 10 – Whole-brain spatial adjacency binary matrix, the ROIs are ordered as in section 2.

| | Anesthetized female | | | Anesthetized male | | | Awake male | | |
|---|---|---|---|---|---|---|---|---|---|
| Model predictors | $\beta$ | $R^2$ | AIC | $\beta$ | $R^2$ | AIC | $\beta$ | $R^2$ | AIC |
| **CO** | 0.645 | 0.416 | 6911 | 0.641 | 0.411 | 6934 | 0.612 | 0.375 | 7118 |
| **VS** | 0.398 | 0.159 | 8007 | 0.389 | 0.151 | 8034 | 0.292 | 0.086 | 8260 |
| **CGE** | 0.557 | 0.310 | 7410 | 0.555 | 0.308 | 7421 | 0.495 | 0.245 | 7683 |
| **ED** | 0.515 | 0.265 | 7600 | 0.538 | 0.290 | 7499 | 0.630 | 0.397 | 7009 |
| **SA** | 0.550 | 0.302 | 7446 | 0.575 | 0.330 | 7322 | 0.609 | 0.371 | 7136 |
| **ED**, **SA** | 0.289 | 0.355 | 7210 | 0.301 | 0.388 | 7054 | 0.413 | 0.479 | 6571 |
| | 0.375 | | | 0.393 | | | 0.360 | | |
| **CO**, **VS** | 0.583 | 0.479 | 6572 | 0.581 | 0.470 | 6624 | 0.576 | 0.397 | 7012 |
| | 0.258 | | | 0.248 | | | 0.153 | | |
| **CO**, **ED**, **SA** | 0.452 | 0.464 | 6657 | 0.411 | 0.478 | 6578 | 0.262 | 0.516 | 6355 |
| | 0.097 | | | 0.126 | | | 0.302 | | |
| | 0.223 | | | 0.255 | | | 0.272 | | |
| **VS**, **ED**, **SA** | 0.291 | 0.436 | 6811 | 0.275 | 0.460 | 6679 | 0.164 | 0.505 | 6422 |
| | 0.281 | | | 0.293 | | | 0.409 | | |
| | 0.319 | | | 0.340 | | | 0.328 | | |
| **CO**, **CGE**, **ED**, **SA** | 0.389 | 0.493 | 6493 | 0.355 | 0.502 | 6441 | 0.249 | 0.517 | 6350 |
| | 0.227 | | | 0.205 | | | 0.047 | | |
| | 0.015 | | | 0.053 | | | 0.286 | | |
| | 0.196 | | | 0.230 | | | 0.267 | | |
| **CO**, **VS**, **ED**, **SA** | 0.402 | 0.520 | 6329 | 0.363 | 0.529 | 6272 | 0.234 | 0.533 | 6247 |
| | 0.245 | | | 0.233 | | | 0.137 | | |
| | 0.111 | | | 0.140 | | | 0.310 | | |
| | 0.193 | | | 0.225 | | | 0.255 | | |

TABLE 1 – Functional connectivity models parameters, $R^2$ and AIC using the 78 cerebrum ROIs.

### 2 ROIs used

| Structure | Allen Atlas id |
|---|---|
| Frontal pole, cerebral cortex | 184 |
| Primary motor area | 985 |
| Secondary motor area | 993 |
| Primary somatosensory area, nose | 353 |
| Primary somatosensory area, barrel field | 329 |
| Primary somatosensory area, lower limb | 337 |
| Primary somatosensory area, mouth | 345 |
| Primary somatosensory area, upper limb | 369 |
| Primary somatosensory area, trunk | 361 |
| Supplemental somatosensory area | 378 |
| Gustatory areas | 1057 |
| Visceral area | 677 |
| Dorsal auditory area | 1011 |
| Primary auditory area | 1002 |
| Ventral auditory area | 1018 |
| Anterolateral visual area | 402 |
| Anteromedial visual area | 394 |
| Lateral visual area | 409 |
| Primary visual area | 385 |
| Posterolateral visual area | 425 |
| posteromedial visual area | 533 |
| Anterior cingulate area, dorsal part | 39 |
| Anterior cingulate area, ventral part | 48 |
| Prelimbic area | 972 |
| Infralimbic area | 44 |
| Orbital area, lateral part | 723 |
| Orbital area, medial part | 731 |
| Orbital area, ventrolateral part | 746 |
| Agranular insular area, dorsal part | 104 |
| Agranular insular area, posterior part | 111 |
| Agranular insular area, ventral part | 119 |
| Retrosplenial area, lateral agranular part | 894 |
| Retrosplenial area, dorsal part | 879 |
| Retrosplenial area, ventral part | 886 |
| Posterior parietal association areas | 22 |
| Temporal association areas | 541 |
| Perirhinal area | 922 |
| Ectorhinal area | 895 |
| Main olfactory bulb | 507 |

Continued on next page

| Structure | Allen Atlas id |
|---|---|
| Accessory olfactory bulb | 151 |
| Anterior olfactory nucleus | 159 |
| Taenia tecta | 589 |
| Dorsal peduncular area | 814 |
| Piriform area | 961 |
| Nucleus of the lateral olfactory tract | 619 |
| Cortical amygdalar area, anterior part | 639 |
| Cortical amygdalar area, posterior part | 647 |
| Piriform-amygdalar area | 788 |
| Postpiriform transition area | 566 |
| Field CA1 | 382 |
| Field CA2 | 423 |
| Field CA3 | 463 |
| Dentate gyrus | 726 |
| Entorhinal area, lateral part | 918 |
| Entorhinal area, medial part, dorsal zone | 926 |
| Parasubiculum | 843 |
| Postsubiculum | 1037 |
| Presubiculum | 1084 |
| Claustrum | 583 |
| Endopiriform nucleus, dorsal part | 952 |
| Endopiriform nucleus, ventral part | 966 |
| Lateral amygdalar nucleus | 131 |
| Basolateral amygdalar nucleus | 295 |
| Basomedial amygdalar nucleus | 319 |
| Posterior amygdalar nucleus | 780 |
| Caudoputamen | 672 |
| Nucleus accumbens | 56 |
| Fundus of striatum | 998 |
| Olfactory tubercle | 754 |
| Lateral septal nucleus, caudal (caudodorsal) part | 250 |
| Lateral septal nucleus, rostral (rostroventral) part | 258 |
| Lateral septal nucleus, ventral part | 266 |
| Septofimbrial nucleus | 310 |
| Anterior amygdalar area | 23 |
| Central amygdalar nucleus | 536 |
| Intercalated amygdalar nucleus | 1105 |
| Medial amygdalar nucleus | 403 |
| Globus pallidus, external segment | 1022 |
| Globus pallidus, internal segment | 1031 |
| Substantia innominata | 342 |

Continued on next page

| Structure | Allen Atlas id |
|---|---|
| Magnocellular nucleus | 298 |
| Medial septal nucleus | 564 |
| Diagonal band nucleus | 596 |
| Triangular nucleus of septum | 581 |
| Bed nuclei of the stria terminalis | 351 |
| Ventral anterior-lateral complex of the thalamus | 629 |
| Ventral medial nucleus of the thalamus | 685 |
| Ventral posterolateral nucleus of the thalamus | 718 |
| Ventral posteromedial nucleus of the thalamus | 733 |
| Ventral posteromedial nucleus of the thalamus, parvicellular part | 741 |
| Subparafascicular nucleus, parvicellular part | 422 |
| Peripeduncular nucleus | 1044 |
| Medial geniculate complex, dorsal part | 1072 |
| Medial geniculate complex, ventral part | 1079 |
| Medial geniculate complex, medial part | 1088 |
| Dorsal part of the lateral geniculate complex | 170 |
| Lateral posterior nucleus of the thalamus | 218 |
| Posterior complex of the thalamus | 1020 |
| Posterior limiting nucleus of the thalamus | 1029 |
| Anteroventral nucleus of thalamus | 255 |
| Anteromedial nucleus, dorsal part | 1096 |
| Anteromedial nucleus, ventral part | 1104 |
| Anterodorsal nucleus | 64 |
| Lateral dorsal nucleus of thalamus | 155 |
| Intermediodorsal nucleus of the thalamus | 59 |
| Mediodorsal nucleus of thalamus | 362 |
| Submedial nucleus of the thalamus | 366 |
| Paraventricular nucleus of the thalamus | 149 |
| Parataenial nucleus | 15 |
| Nucleus of reuniens | 181 |
| Rhomboid nucleus | 189 |
| Central medial nucleus of the thalamus | 599 |
| Central lateral nucleus of the thalamus | 575 |
| Reticular nucleus of the thalamus | 262 |
| Ventral part of the lateral geniculate complex | 178 |
| Medial habenula | 483 |
| Lateral habenula | 186 |
| Paraventricular hypothalamic nucleus | 38 |
| Arcuate hypothalamic nucleus | 223 |
| Dorsomedial nucleus of the hypothalamus | 830 |
| Medial preoptic area | 523 |

Continued on next page

| Structure | Allen Atlas id |
|---|---|
| Periventricular hypothalamic nucleus, posterior part | 126 |
| Periventricular hypothalamic nucleus, preoptic part | 133 |
| Subparaventricular zone | 347 |
| Anterior hypothalamic nucleus | 88 |
| Medial mammillary nucleus | 491 |
| Supramammillary nucleus | 525 |
| Medial preoptic nucleus | 515 |
| Ventromedial hypothalamic nucleus | 693 |
| Posterior hypothalamic nucleus | 946 |
| Lateral hypothalamic area | 194 |
| Lateral preoptic area | 226 |
| Subthalamic nucleus | 470 |
| Tuberal nucleus | 614 |
| Superior colliculus, sensory related | 302 |
| Inferior colliculus, central nucleus | 811 |
| Inferior colliculus, dorsal nucleus | 820 |
| Inferior colliculus, external nucleus | 828 |
| Substantia nigra, reticular part | 381 |
| Ventral tegmental area | 749 |
| Midbrain reticular nucleus, retrorubral area | 246 |
| Midbrain reticular nucleus | 128 |
| Superior colliculus, motor related | 294 |
| Periaqueductal gray | 795 |
| Anterior pretectal nucleus | 215 |
| Nucleus of the optic tract | 628 |
| Nucleus of the posterior commissure | 634 |
| Cuneiform nucleus | 616 |
| Red nucleus | 214 |
| Substantia nigra, compact part | 374 |
| Pedunculopontine nucleus | 1052 |
| Interpeduncular nucleus | 100 |
| Central linear nucleus raphe | 591 |
| Nucleus of the lateral lemniscus | 612 |
| Principal sensory nucleus of the trigeminal | 7 |
| Parabrachial nucleus | 867 |
| Superior olivary complex | 398 |
| Pontine central gray | 898 |
| Pontine gray | 931 |
| Pontine reticular nucleus, caudal part | 1093 |
| Supratrigeminal nucleus | 534 |
| Tegmental reticular nucleus | 574 |

Continued on next page

| Structure | Allen Atlas id |
|---|---|
| Motor nucleus of trigeminal | 621 |
| Superior central nucleus raphe | 679 |
| Pontine reticular nucleus | 146 |
| Dorsal cochlear nucleus | 96 |
| Ventral cochlear nucleus | 101 |
| Nucleus of the solitary tract | 651 |
| Spinal nucleus of the trigeminal, caudal part | 429 |
| Spinal nucleus of the trigeminal, interpolar part | 437 |
| Spinal nucleus of the trigeminal, oral part | 445 |
| Facial motor nucleus | 661 |
| Gigantocellular reticular nucleus | 1048 |
| Inferior olivary complex | 83 |
| Intermediate reticular nucleus | 136 |
| Lateral reticular nucleus | 235 |
| Magnocellular reticular nucleus | 307 |
| Medullary reticular nucleus, dorsal part | 1098 |
| Medullary reticular nucleus, ventral part | 1107 |
| Parvicellular reticular nucleus | 852 |
| Paragigantocellular reticular nucleus, dorsal part | 970 |
| Paragigantocellular reticular nucleus, lateral part | 978 |
| Nucleus prepositus | 169 |
| Lateral vestibular nucleus | 209 |
| Medial vestibular nucleus | 202 |
| Spinal vestibular nucleus | 225 |
| Superior vestibular nucleus | 217 |
| Hypoglossal nucleus | 773 |
| Central lobule | 920 |
| Culmen | 928 |
| Pyramus (VIII) | 951 |
| Nodulus (X) | 968 |
| Simple lobule | 1007 |
| Ansiform lobule | 1017 |
| Paramedian lobule | 1025 |
| Paraflocculus | 1041 |
| Flocculus | 1049 |
| Fastigial nucleus | 989 |
| Interposed nucleus | 91 |
| Dentate nucleus | 846 |
